# Divergent Macrophage-Regulated T cell States Determine Response to Bacillus Calmette-Guérin (BCG) in High-Risk Bladder Cancer

**DOI:** 10.1101/2025.09.18.677132

**Authors:** Ryan J. Brown, Mairah T. Khan, Hongshen Niu, Joseph R. Podojil, Bonnie Choy, Weiguo Cui, Joshua J. Meeks

## Abstract

The primary therapy for high-risk bladder cancer (BCa) is repeated instillations of the tuberculosis vaccine, Bacillus Calmette-Guerin (BCG). While BCG decreases the risk of recurrence by more than half, the concerted mechanisms of immune activation from BCG are unknown. Our objective was to investigate how the immune response differs between responders and non-responders to BCG therapy. We performed single-cell RNA-sequencing of isolated immune cells adjacent to high-risk bladders before and after BCG in BCG responders and non-responders. We identify an increase in Th17-like Th1 cells in BCG responders, characterized by greater expression of pro-inflammatory cytokines. Alternatively, non-responders had increased CD8+ T-cell exhaustion and T-regulatory cells. We identify that the primary mechanism of divergent T cell activity is driven by altered polarization and immunosuppressive signaling with myeloid cells. Through a machine-learning-based approach, we identified a Th17-like Th1 cytokines, such as IL17, IL21, and IL26, were predictive of a response, which were then validated in a separate BCG-treated BCa cohort. Together, this suggests that dynamic regulation of myeloid-T cell interactions can be targeted to improve BCG activity.

## Introduction

Bladder cancer (BCa) is the eighth most common cancer worldwide and the fourth most common among men^1–3^. It is also the most economically burdensome malignancy, largely due to its high recurrence rate, which necessitates frequent and invasive monitoring procedures such as cystoscopies to detect new or recurrent tumors^4^. Most cases (approximately 80%) are confined to the superficial lining of the bladder and are classified as non-muscle invasive bladder cancer (NMIBC)^1,3^. Standard management of NMIBC involves surgical resection followed by adjuvant immunotherapy, but nearly a third of tumors recur within two years^5^. The most effective and widely used therapy is intravesical administration of *Mycobacterium bovis* Bacillus Calmette-Guérin (BCG), which has reduced recurrence by more than 50%^6^. However, despite its clinical utility, approximately one-third of patients fail to respond to BCG^7–9^. Among these BCG-unresponsive cases, 25–30% progress to muscle-invasive disease, necessitating systemic therapy or radical cystectomy^10^. Compounding these challenges, global manufacturing shortages have further constrained access to BCG, prompting rationing strategies in many regions^11^.

A major limitation to developing new strategies to improve or replace BCG is the limited knowledge of the immune mechanisms by which BCG causes BCa eradication. Initially developed as a latent vaccine to prevent infection with *mycobacterium tuberculosi*s (*Mtb*), the administration of BCG was identified to have therapeutic activity in multiple solid tumors^12^. Extensive immune characterization of M. tuberculosis (*Mtb)* has identified several processes that may be critical to BCG activity. *Mtb* infects the alveolar macrophages of the lung, resulting in activation of the innate immune system and recruitment of Th1 CD4+ T cells that ultimately create granulomas and tissue necrosis^13^. Conversely, *Mtb* dampens the immune response through induction of immunosuppressive cells and signaling such as TGFb and IL10, the downregulation of MHCII on antigen-presenting cells, and latent pulmonary tuberculosis has been associated with exhaustion in CD4+ and CD8+ T cells^14–16^. Attempts to ameliorate exhaustion through PD1 immune checkpoint immunotherapy and T cell exhaustion in BCa remains controversial, but tumors with increased expression of PDL1 have been reported to have diminished response to BCG^17^. Pembrolizumab was approved for BCG-unresponsive BCa in 2019, and a recent clinical trial combining anti-PD1 therapy with BCG prolongs recurrence of BCa^18,19^. Therefore, investigating the mechanisms of immune-mediated BCG could help identify how to synergize immune checkpoint blockade with BCG and discover new therapies for treating patients who are unresponsive or unable to receive adequate BCG.

To evaluate how the tumor microenvironment of the bladder evolves during BCG therapy, we performed single-cell transcriptome analysis of the adjacent normal bladder tissue and circulating immune cell populations from patients treated with BCG for high-risk bladder cancer (BCa). Through cellular characterization, cell-cell interactions, and machine learning, we compare BCG-naïve to BCG-responsive and non-responsive bladders, identifying immune populations and gene expression programs that parallel changes found in latent *M. Tuberculosis.* Specifically, we identify a Th17-like Th1 CD4+ T cell population that associates with clinical response to BCG. Interestingly, we found that BCG-responders had a more activated macrophage profile with an increase in antigen presentation. Conversely, BCG non-responder macrophages exhibited reduced MHC II presentation, increased TGF-β signaling, and enhanced co-inhibitory pathways. In alignment with this, BCG nonresponding patients had a corresponding increase in immunosuppressive Tregs and exhausted CD8+ T cells. Together, these results suggest that strategies applied to activate the immune system in *Mtb* and potentially blockade of exhaustion could be leveraged to overcome BCG resistance in patients with BCa.

## Results

To investigate how the local and systemic immune systems evolve during intravesical BCG treatment for bladder cancer, single-cell immune profiling (single-cell RNA-Seq, scRNA-Seq) was performed on blood and tissue biopsies from patients with high-risk non-muscle-invasive bladder cancer (Figure 1A). To avoid possible contribution and over-representation of epithelial cells from invasive BCa, tissue biopsies were taken adjacent to the tumor in visually “normal” areas of the bladder. CD45 enrichment was performed on the tissue samples during the processing before scRNA-Seq (Figure 1A). To further elucidate the mechanism of an effective BCG response, data were collected from BCG-naïve patients (n = 7), BCG-responsive patients (n = 6), and BCG-unresponsive patients (n = 8) among a total of 21 patients (Figure 1B-C). In parallel, scRNA-Seq was performed on PBMCs to augment the analysis of tissue specimens (Figure 1A-C).

**Figure 1.**
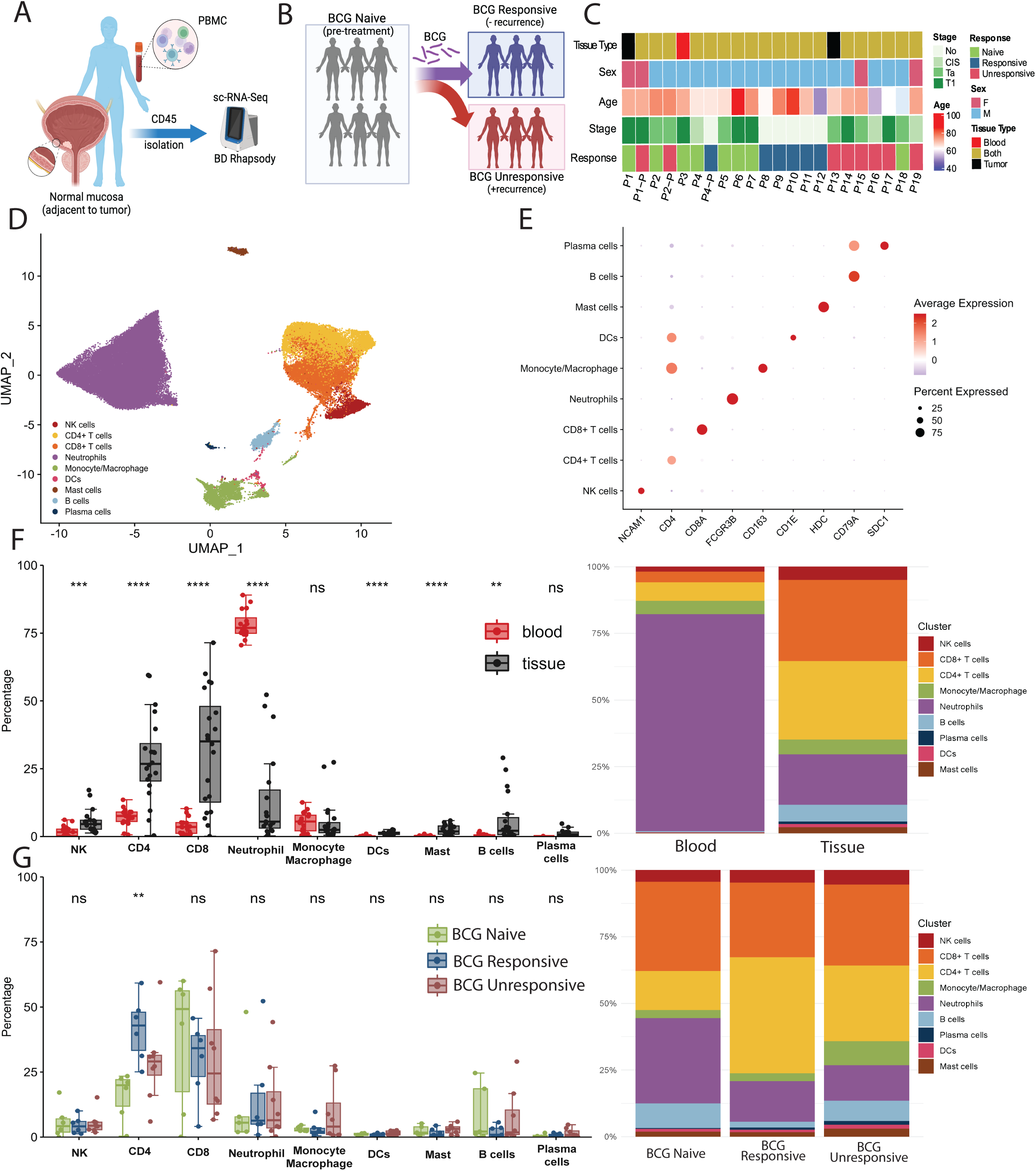
Comprehensive dissection and clustering of immune cells from BCG-treated bladders. **a.** Sample collection from patients with bladder cancer. For most patients, whole and bladder biopsies adjacent to tumor were collected prospectively with CD45+ isolation and analysis by single-cell RNA-Seq using the BD Rhapsody system **b.** Patients were prospectively enrolled at three distinct time points: BCG naïve (prior to BCG therapy), and post BCG. After BCG therapy, patients were classified as either responsive or resistant, depending on the result of the bladder biopsy. **c.** Documentation of each patient’s covariates including sex, age, and tumor stage as well as documentation of which tissue compartments were sequenced and the patient’s BCG response. **d.** UMAP plot of all immune cells collected across 19 patients from the blood and bladder tissue from BCG naïve, BCG responders, and BCG non-responders. **e.** Dot plot of representative genes expressed in each major cluster. Dot size represents percent of cells expressing the gene; color represents scaled expression of the gene. **f.** Proportion of immune cell types in blood and bladder. Left: Box plots showing the distribution of immune cell type proportions for each patient. Right: Stacked bar plots summarizing immune cell type proportions after down sampling to 200 cells per sample and measured by a chi-square test. **g.** Proportion of immune cell types in BCG naïve, responsive, and unresponsive groups in the bladder tissue. Left: Box plots showing the distribution of immune cell type proportions. Right: Stacked bar plots summarizing immune cell type proportions after down sampling to 200 cells per sample and measured by a chi-square test. Box plots (median, interquartile range, and whiskers extending to 1.5 × IQR) in f and g. * P < 0.05, ** P < 0.01, *** P < 0.001, **** P < 0.0001.

### Immune landscape of the bladder and circulation of BCG-treated patients

scRNA-Seq was generated for 84,616 cells using the BD Rhapsody single-cell system, with matched tissue and blood samples multiplexed together using sample tags. The BD system was selected to allow for the preservation of fragile neutrophil populations from the bladder^20^. After quality control analyses, data were analyzed for 74,651 immune cells. We utilized the R package Seurat to integrate the data from 20 blood and 21 bladder biopsies across BCG naïve, BCG responder, and BCG non-responder groups using canonical correlation analysis. After integrating the samples, we projected the immune cells in two dimensions by Uniform Manifold Approximation and Projection (UMAP) and performed unsupervised clustering (Figure 1D). We identified immune cell populations by using differentially expressed genes across these identified clusters (Figure 1E, Suppl. 1A). To confirm our sequencing data quality, we plotted cell counts and the distribution of cell types for each patient in the tissue (Suppl. 2A) and blood (Suppl. 2B). We next examined how immune cell composition varied between the peripheral blood and the tumor site (Figure 1 F-G, Suppl. 1B). A global comparison of all CD45_⁺_ cells revealed striking compartment-specific differences of immune cells between bladder tissue (n=32,873) and blood (n=41,778). We identified that blood samples were predominantly comprised of neutrophils (81% of cells) while T cells were the primary CD45+ cells isolated from the bladder (58% of cells) (Figure 1F). Underscoring the distinct immune landscape at the tumor site, the bladder tissues exhibited significantly higher frequencies of NK cells, CD4+ and CD8+ T cells, dendritic cells, mast cells, and B cells compared to blood (Figure 1F). To understand how BCG influences the immune profile, we categorized samples based on BCG exposure status (pre-BCG [naïve] and post-BCG [BCG-responsive or BCG-unresponsive]) and compared the relative frequencies of conditions across immune cell types (Figure 1G). In the bladder, BCG exposure was linked to a notable increase in CD4+ T cells (Figure. 1G), while the peripheral blood showed no significant change in overall immune composition (Suppl. 1C). However, this increase in CD4+ T cells was mainly driven by BCG-responsive tumors, indicating a localized, treatment-related expansion of this subset in effective immune responses (Figure 1G).

### Opposing Th17 and Treg signatures define response to BCG

Because they represent the greatest abundance of immune cells in the bladder, we first analyzed the lymphocyte pool including CD4+ (Figure 2), CD8+ (Figure 3), and NK cells (Suppl. Figure 3). We further examined the CD4+ T cell landscape before and after BCG administration since we identified a significant expansion of CD4+ T cells in the bladders of BCG-responsive patients. Overall, there were 11,757 CD4+ T cells across all patients and cellular compartments. To more accurately align the CD4+ T cells, we performed integration using SCTransform, followed by a UMAP projection and nearest neighbor clustering (Figure 2A). To identify cell subtypes, we performed differential gene expression and identified previously described canonical CD4+ T cell types. The cluster with high expression of *TCF7*, *SELL*, *LEF1,* and *CCR7* was identified as “naïve/memory cluster”, while the cluster with high expression of *FOXP3*, *IL2RA,* and *IKZF2* was identified as “Tregs” (Figure 2B). Most other CD4+ T cells exhibited a Th1 phenotype, characterized by high levels of *STAT4, IL-12R*β*2,* and *IFN-*γ. However, some of these cells exhibited a more conventional Th1 profile, characterized by high expression of *TNF, TBX21,* and *NKG7*, while others displayed a unique expression of Th17-like markers, including *IL-17A, IL-17F, SOX5,* and *IL-26* (Figure 2B). To further investigate the differences between these CD4+ T cells, we applied a cosine similarity index, which performs pairwise comparisons of gene expression between cell types. We found that Tregs and naïve/memory CD4+ T cells had little similarity to all other CD4+ cell subtypes, while Th1 and Th17-like Th1 cells had high pairwise cosine similarity scores, suggesting that these Th17-like cells are still part of the Th1 lineage rather than fully differentiated Th17 cells (Suppl. 4A).

**Figure 2.**
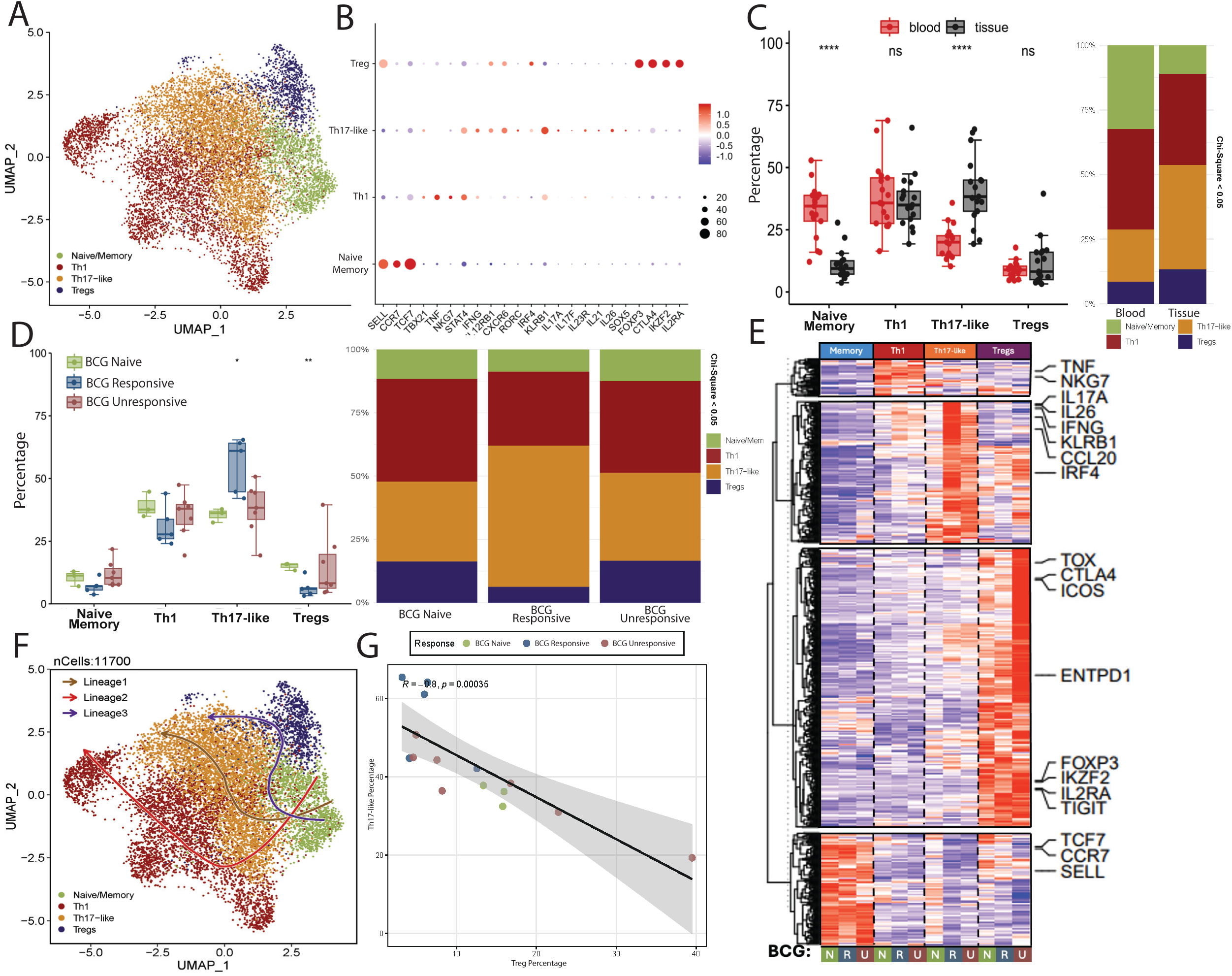
Opposing Th17 and Treg signatures define response to BCG. **a.** UMAP plot of CD4+ T cells from the blood and bladder tissue from BCG naïve, BCG responders, and BCG non-responders. **b.** Dot plot of representative genes expressed in each major cluster. Dot size represents percent of cells expressing the gene; color represents scaled expression of the gene. **c.** Proportion of CD4+ cell subsets in blood and bladder. Left: Box plots showing the distribution of immune cell type proportions for each patient. Right: Stacked bar plots summarizing immune cell type proportions after down sampling to 100 cells per sample. **d.** Proportion CD4+ cell subsets in BCG naïve, responsive, and unresponsive groups in the bladder tissue. Left: Box plots showing the distribution of CD4+ cell subset proportions. Right: Stacked bar plots summarizing CD4+ cell subset proportions after down sampling to 100 cells per sample. **e.** Heatmap showing differentially expressed genes of CD4+ T cell subsets categorized by BCG naïve, responsive and unresponsive groups. **i.** Slingshot trajectory plot showing predicted cellular differentiation possibilities of CD4+ T cells. **j**. Linear regression between a patient’s proportion of Th17-like and Treg cells. Each point represents one patient. Gray shaded region, 95% confidence interval. Box plots (median, interquartile range, and whiskers extending to 1.5 × IQR) in c and d. * P < 0.05, ** P < 0.01, *** P < 0.001, **** P < 0.0001.

**Figure 3.**
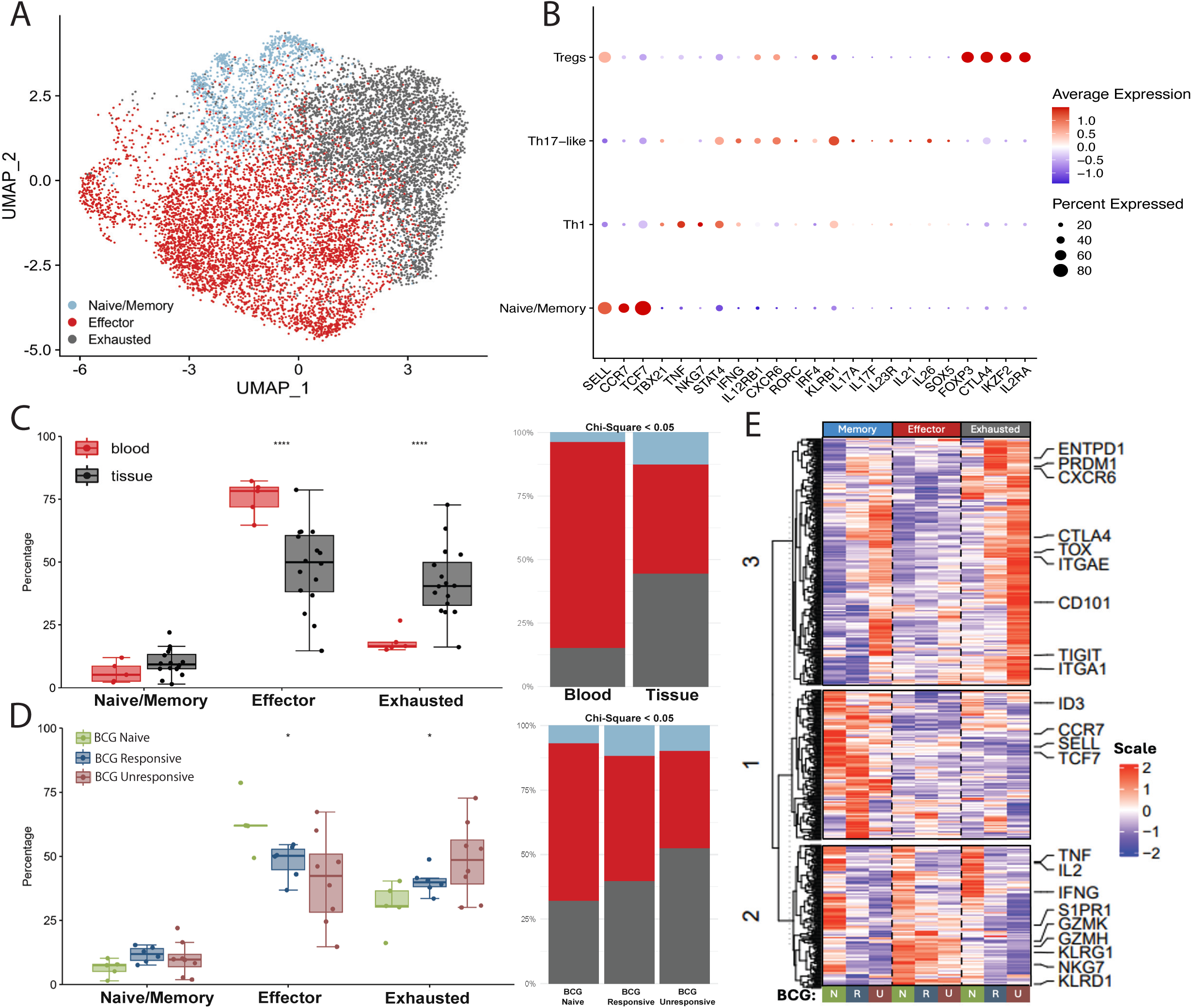
CD8+ T cells demonstrate increased exhaustion in BCG-unresponsive bladders. **a.** UMAP plot of CD8+ T cells from the blood and bladder tissue from BCG naïve, BCG responders, and BCG non-responders. **b.** Dot plot of representative genes expressed in each major cluster. Dot size represents percent of cells expressing the gene; color represents scaled expression of the gene. **c.** Proportion of CD8+ cell subsets in blood and bladder. Left: Box plots showing the distribution of immune cell type proportions for each patient. Right: Stacked bar plots summarizing CD8+ subset proportions after down sampling to 200 cells per sample. **d.** Proportion CD8+ cell subsets in BCG naïve, responsive, and unresponsive groups in the bladder tissue. Left: Box plots showing the distribution of CD8+ T cell subset proportions. Right: Stacked bar plots summarizing CD8+ cell subset proportions after down sampling to 200 cells per sample. **e.** Heatmap showing differentially expressed genes of CD8+ T cell subsets categorized by BCG naïve, responsive and unresponsive groups. Box plots (median, interquartile range, and whiskers extending to 1.5 × IQR) in c and d. * P < 0.05, ** P < 0.01, *** P < 0.001, **** P < 0.0001.

To analyze the immune profile of CD4+ T cells during bladder cancer treatment with BCG, we compared the cell types between blood and bladder tissue (Suppl. 4B). While the proportion of Th1 cells was consistent between treated and untreated bladder samples (∼40%), Th17-like Th1 cells were markedly enriched in the bladder, emerging as the dominant CD4+ population (∼45%) (Figure 2C). This shift was accompanied by a significant depletion of naïve/memory CD4+ T cells, reflecting a transition toward a more activated immune environment in the bladder (Figure 2C). We next determined how BCG exposure status impacted CD4+ T cell distributions. While BCG exposure had minimal impact on circulating immune cell populations (Suppl. 4C), we found that BCG-responsive patients had increased Th17-like Th1 cells and decreased Tregs when compared to BCG naïve patients (Figure 2D). While Th1 changes have been described after BCG exposure, the Th17-like signature in BCG-responsive patients is consistent with the activation of a Th17 CD4+ T cell population after *Mtb* exposure^21^.

To more narrowly explore how BCG exposure and response impact tissue-specific CD4+ T cells after BCG exposure, we investigated how gene expression was altered within these identified cell states. We identified the top differentially expressed genes and performed down-sampling to ensure each patient was weighted equally. We then applied k-means hierarchical clustering to identify broad gene expression signatures. We identified four major gene signatures corresponding to each of the four major cell types identified in our UMAP clustering (Figure 2E). While many cell-type-specific genes were expressed uniquely in each cluster, we also observed differences in gene expression between BCG exposures. Interestingly, we observed that the BCG-responsive group exhibited high gene expression of Th17 genes, such as *IL17A* and *IL26*, as well as higher expression of Th1 genes, including *IFNG,* within the Th17-like Th1 cell cluster. Together, this suggests that the Th17-like Th1 population is not only more frequent in BCG responders but also has a greater capacity to produce critical functional markers, indicating greater activation and function on a per-cell basis. In contrast, nonresponding BCG patients had higher expression of Treg signature genes, including *CTLA4, ICOS, FOXP3,* and *TOX.* Higher expression of these immunosuppressive surface markers, cytokines, and transcription factors suggests that BCG-unresponsive patients have Tregs with more potent immunosuppressive capacity. Th17 cells and Tregs have been previously shown to have mutual antagonism, competing for TGF-β during differentiation and subsequently establishing opposing inflammatory environments, a phenomenon well established in autoimmune diseases^22^. This finding aligns with past work that identified a relationship between CD4+ T cells and bladder cancer, whereby the percentage of IL-17-producing CD4+ T cells inversely correlate with the percentage of Treg cells^23^. To explore this dynamic in our BCG bladder cancer model, we performed trajectory analysis with the slingshot algorithm and identified three distinct lineages (Figure 2F). We observed that these lineages originated from the naïve/memory CD4+ T cell population and differentiated into the Th1, Th1/17, and Treg populations. Overall, our model suggests an early branch point prior to terminal fate commitments. Consistent with the mutual antagonism model of differentiation, we observed a negative correlation between Th1/17 and Treg populations across samples (Figure 2G). Together, our results suggest that this Th17/Treg relationship is critical for generating effective immune responses to mycobacterial infections such as BCG and *Mtb*.

### CD8+ T cells demonstrate increased exhaustion in BCG-unresponsive bladders

While there were no significant differences in the overall proportion of CD8+ T cells between responders and non-responders to BCG (Figures 1H and I), we sought to identify differences in CD8+ T cell subsets using single-cell profiling. We isolated the CD8+ T cell population, reintegrated the cells, and applied unbiased UMAP clustering, where we identified three distinct clusters through shared nearest neighbor optimization (Figure. 3A). These CD8+ subtypes had varied cell expressions that reflected previously described cell types including “naïve/memory T cells,” (*TCF7, KLF2, SELL and CCR7*), “effector T cells,” (*KLRG1, PRF1, and CXC3R1*), and “exhausted T cells” (*TOX, ENTPD1, LAG3*) (Figure 3B, Suppl. 5A)^24,25^. To explore how bladder tissue affects the CD8+ T cell populations, we compared the shifts in population between the blood and bladder. We observed a higher number of effector CD8+ T cells in the blood, and an enrichment of exhausted CD8+ T cells in the bladder (Figure. 3C, Suppl. 5B). This tissue-specific shift in CD8+ T cell profile suggests bladder cancer promotes localized T cell exhaustion.

To determine if BCG-responsive and unresponsive patients had differences within CD8+ T cells, we compared the frequency of CD8+ cell subtypes. While there was no difference in CD8+ cell types in the blood (Suppl. 5C), we identified significantly lower effector CD8+ T cells in BCG unresponsive bladders and significantly increased exhausted CD8+ T cells in unresponsive bladders (Figure 3D). To more narrowly explore how BCG exposure impacts CD8+ T cell differentiation, we analyzed the differential expression of individual genes within each cell across both CD8+ clusters and BCG exposures, identifying a total of 1,064 genes. To control differences in patient cell count, each patient sample was randomly down-sampled to 200 cells to ensure equal weighting between patient replicates. We then applied k-means hierarchical clustering so that we could identify broad gene expression signatures (Figure 3E). We found that gene expression could be categorized into three main signatures, including a Tpro, Teff, and Texh gene signature. While these signatures corresponded to CD8+ T cell clusters, the signatures were not equally shared across BCG exposures within those clusters. In the naïve/memory cluster, the Tpro signature was more highly expressed in the BCG naïve and BCG responders, while the BCG non-responders indicated a lower level of expression. Notably, recent investigations of CD8+ T cells have highlighted the significance of these stem-like signatures in sustaining robust CD8+ T cell responses against tumors^26,27^. The Teff gene cluster exhibited a shared signature of effector-related genes, including *GZMK, GZMH, KLRG1, S1PR1,* and *NKG7,* across the BCG exposures.

However, a second effector signature comprised of cytokines such as *TNF, IL2,* and *IFNG* was specific to the BCG naïve condition, with a significant reduction in patients exposed to BCG. Finally, we identified a third signature made up of genes related to exhaustion. Overall, we found that some genes were shared across BCG exposures in the exhausted cluster, including *ENTPD1, CXCR6,* and *PRDM1.* However, the BCG non-responding patients showed significantly higher expression of additional exhaustion markers, including *CTLA4, TOX,* and *TIGIT,* as well as tissue-residence-related markers, including *ITGAE, CD101,* and *ITGA1.* Notably, we found that this exhausted CD8+ T cell signature was also higher in the naïve/memory cluster, indicating that these precursor cells may already be undergoing exhaustion. Interestingly, the anti-PD-1 therapy function is known to operate by inhibiting PD-1 on these progenitor cells^28^. These results suggest that corresponding immunotherapies may be beneficial for BCG nonresponding patients who exhibit high levels of CD8+ T cell exhaustion. Overall, this analysis reveals that BCG non-responding patients exhibit altered CD8+ T cell differentiation and cellular signatures characteristic of T-cell exhaustion.

### Macrophage polarization and T cell interactions

We further evaluated the myeloid populations, including macrophages (Figure 4) and neutrophils (Suppl. Figure 6). *Mtb* primarily infects alveolar macrophages in the lung, and numerous studies have implicated macrophage-mediated signaling as a central mechanism of tuberculosis-driven immune regulation^29,30^. Given the parallels between *Mtb* infection and intravesical BCG therapy, we sought to investigate how BCG influences the myeloid compartment within the bladder. To investigate this, we re-integrated all monocyte and macrophage cells, applied UMAP dimensionality reduction, and performed nearest-neighbor clustering, which revealed four major clusters (Figure 4A). Myeloid cell populations were primarily composed of classical monocytes (*CD14, VCAN*) and non-classical monocytes (*FCGR3A, CX3CR1, ITGAL*) (Figure 4B). Some cells shared the expression signature of classical monocytes but also had a strong interferon-stimulated signature (*ISG15, ISG20*) (Figure 4B). Finally, a cluster of macrophages with a primarily M2-like phenotype was identified (*CD163, MRC1*), which we termed tumor-associated macrophages (TAMs) (Figure 4B). Having defined these distinct myeloid subsets, we sought to understand how their distribution differed between systemic circulation and the bladder microenvironment (Suppl. Figure 7A). Reflecting the capacity of tissue environments to drive monocyte-to-macrophage differentiation, we found that bladder tissue exhibited a reduction in circulating monocyte populations and a notable expansion of macrophages (Figure 4C). To explore how BCG influences the myeloid compartment, we compared myeloid populations between BCG-naïve and BCG-exposed patients. Interestingly, classical monocytes, which are primarily involved in inflammatory responses, were increased in the blood of BCG-responsive patients (Figure 4D, Suppl. 7B)^31,32^. Overall, this shift suggests that BCG responders induce a systemic bias toward an inflammatory monocyte phenotype, which in turn modulates the tumor microenvironment upon tissue infiltration and macrophage differentiation.

**Figure 4.**
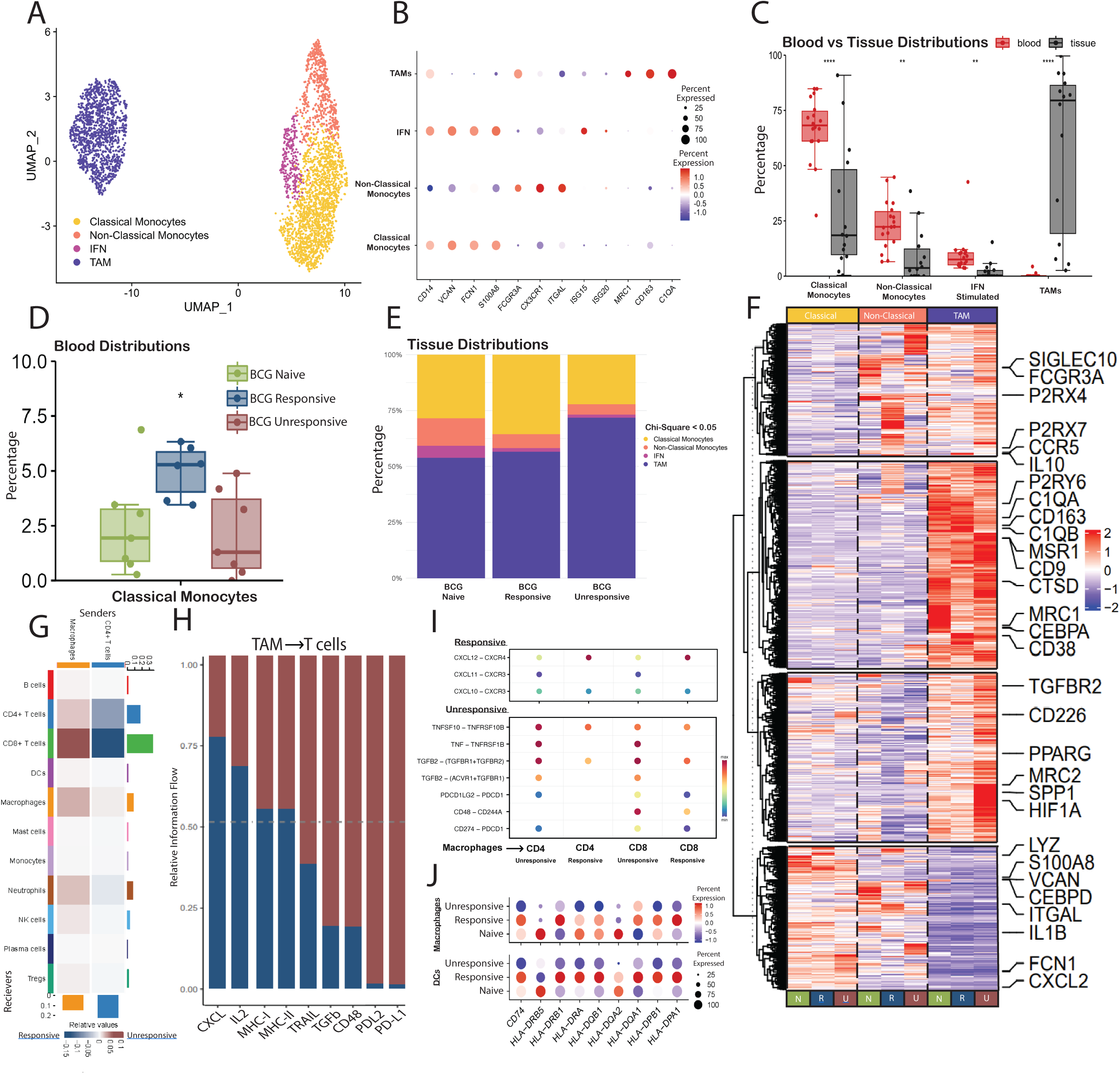
Macrophage Polarization and T cell Interactions. **a.** UMAP plot of monocyte and macrophage cells from the blood and bladder tissue from BCG naïve, BCG responders, and BCG non-responders. **b.** Dot plot of representative genes expressed in each major cluster. Dot size represents percent of cells expressing the gene; color represents scaled expression of the gene. **c.** Proportion of monocyte and macrophage subsets between the blood and bladder. Each dot represents a patient. **d.** Proportion monocyte and macrophage in BCG naïve and BCG exposed (responsive and unresponsive) groups in the blood. **e.** Stacked bar plots summarizing monocyte and macrophage cell subset proportions after down sampling to 50 cells per sample in the bladder tissue. Significance measured by chi-squared test. **f.** Heatmap showing differentially expressed genes of monocyte and macrophage subsets categorized by BCG naïve, responsive and unresponsive groups. **g.** Heatmap of CellChat’s cell-cell interactions scores between BCG responsive and BCG unresponsive cells. **h.** Differences in tumor associated macrophages to T cell interaction scores between BCG responsive and BCG unresponsive cells. **i.** Dot plot of macrophage ligands/cytokines to T cell receptors that are increased (lower) and decreased (upper) in BCG non-responders. Color represents communication probability of a ligand-receptor signaling pathway. **J**. Dot plot of MHCII genes in antigen presenting cells. Dot size represents percent of cells expressing the gene; color represents scaled expression of the gene. Box plots (median, interquartile range, and whiskers extending to 1.5 × IQR) in c and d. * P < 0.05, ** P < 0.01, *** P < 0.001, **** P < 0.0001.

### To explore how BCG impacts macrophage differentiation in the bladder, we examined overall macrophage populations

We found that the majority (1,071 cells) of the total macrophages (1,284 cells) were present in the BCG-unresponsive bladders, with a limited number of macrophages (213 cells, 17%) found in the BCG-responsive bladders. While we saw a trend towards greater TAMs across BCG conditions within the bladder tissue, there was a high degree of heterogeneity across patients resulting in no significant differences across groups (Suppl. 7C). However, when we performed down-sampling and quantified cellular differences, we did see that BCG-unresponsive patients had the highest proportion of TAMs (Figure 4E). TAMs have been described as having both pro-inflammatory and immunosuppressive roles. Although we did not observe discrete subsets corresponding to classically defined M1 and M2 macrophages, we hypothesized that the overall macrophage compartment could still exhibit phenotypic polarization between BCG responders and non-responders. To test this, we downsampled to control for inter-patient differences in cell numbers and performed differential gene expression analysis between cells from BCG-naïve, BCG-responsive, and BCG-unresponsive patients (Figure 4F). Notably, TAMs from BCG non-responders had elevated expression of a distinct cluster of genes such as *SPP1*, which has been implicated in regulation of T cell exhaustion, as well as *PPARG*, *TGFBR2* and *MRC2*, which are associated with regulatory or immunosuppressive functions^33,34^.

To better understand whether TAMs orchestrate immune responses to BCG, we utilized the CellChat algorithm to analyze ligand–receptor interactions across various immune cell types. By weighing the total outgoing signaling from each population and comparing BCG responders to non-responders, we observed notable shifts in cellular crosstalk (Figure 4G). While overall cell–cell interaction strength was largely comparable between groups, we identified striking differences in signaling directed toward CD8+ T cells. In BCG-unresponsive patients, CD8+ T cells received predominant input from TAMs, whereas in BCG-responsive patients, CD4+ T cells were the main signaling source (Figure 4G). This pattern aligns with recent evidence that CD4+ T cell help is essential for sustaining CD8+ T cell function and preventing exhaustion, while TAMs are implicated in regulation of CD8+ T cell dysfunction in BCG unresponsive bladders^24^.

To dissect the pathways mediating these interactions, we compared CellChat signaling profiles between groups. PD-L1 and PD-L2 signaling were the most enriched inhibitory pathways in BCG-unresponsive patients, accompanied by elevated TIGIT, CD244A, and CD266 signaling, as well as immunosuppressive cytokines including TNFSF10 (TRAIL) and TGF-β (Figures 4H-I). In contrast, BCG-responsive patients exhibited increased signaling through pro-inflammatory and recruitment pathways, including CXCL9, CXCL10, CXCL11, CXCL12, and CXCL16, along with enhanced IL-2 and MHC class II pathways (Figure 4H-I). Interestingly, BCG is known to suppress MHC expression^35^, and MHCII expression was not exclusive to BCG responding TAMs, but was more highly expressed across all BCG responding antigen-presenting cells (Figure 4J, Suppl. 8A). To validate this finding, we analyzed our previously published bulk RNA-seq dataset from 103 BCG-treated Stage I NMIBC tumors^36^. Higher expression of antigen presentation genes such as *CD74* and *HLA-DOA* was associated with significantly improved survival, supporting the importance of high MHC signaling for effective responses (Suppl. 8B).

To further investigate how CD4+ T cells may support CD8+ T cell function in BCG responders, we specifically examined signaling between these two populations. While no signaling pathways were strongly enriched in unresponsive patients, we identified a significant enrichment of the CCL5 (RANTES)–CCR5 axis in BCG responders (Suppl. 8C). Although IL-21 signaling is widely recognized as a central mediator of CD4+ T cells to CD8+ T cells, recent studies have shown that RANTES also contributes to the differentiation of CD8+ T cells into the effector lineage^37^.

### Machine learning model to infer important cellular and gene characteristics of BCG responders

Having constructed a single-cell atlas of the immune landscape in bladder cancer, we next aimed to quantify the relative contribution of each immune cell type in predicting BCG responsiveness. We applied the Precise (Predictive Response Analysis from Single-Cell Expression) machine learning framework, which leverages the XGBoost algorithm in a leave-one-out cross-validation setup^38^. Feature importance was refined using Boruta selection, ultimately identifying 57 highly informative genes (Suppl. 9A). To further dissect the impact of these genes on model performance, we calculated SHAP (SHapley Additive exPlanations) values, which quantify each gene’s contribution toward predicting BCG response (Suppl. 9B). Notably, CD4+ T cell–derived cytokines such as IL17A, IL21, and IL26, along with the receptor IL12RB2, were among the strongest positive predictors of response (Figure 5A). These findings align with our earlier observations, which implicated Th17-like Th1 cells in effective immune responses. Moreover, expression of *CD74*, a key component of antigen presentation by antigen presenting cells (APCs), also positively influenced response prediction (Figure 5A, Suppl. 9B). In contrast, genes enriched in suppressive myeloid populations, such as *SPP1* in tumor-associated macrophages, were predictive of BCG non-responsiveness (Figure 5A). Together, this integrative analysis highlights both molecular signatures that distinguish responders from non-responders and provides a framework for future biomarker development.

**Figure 5.**
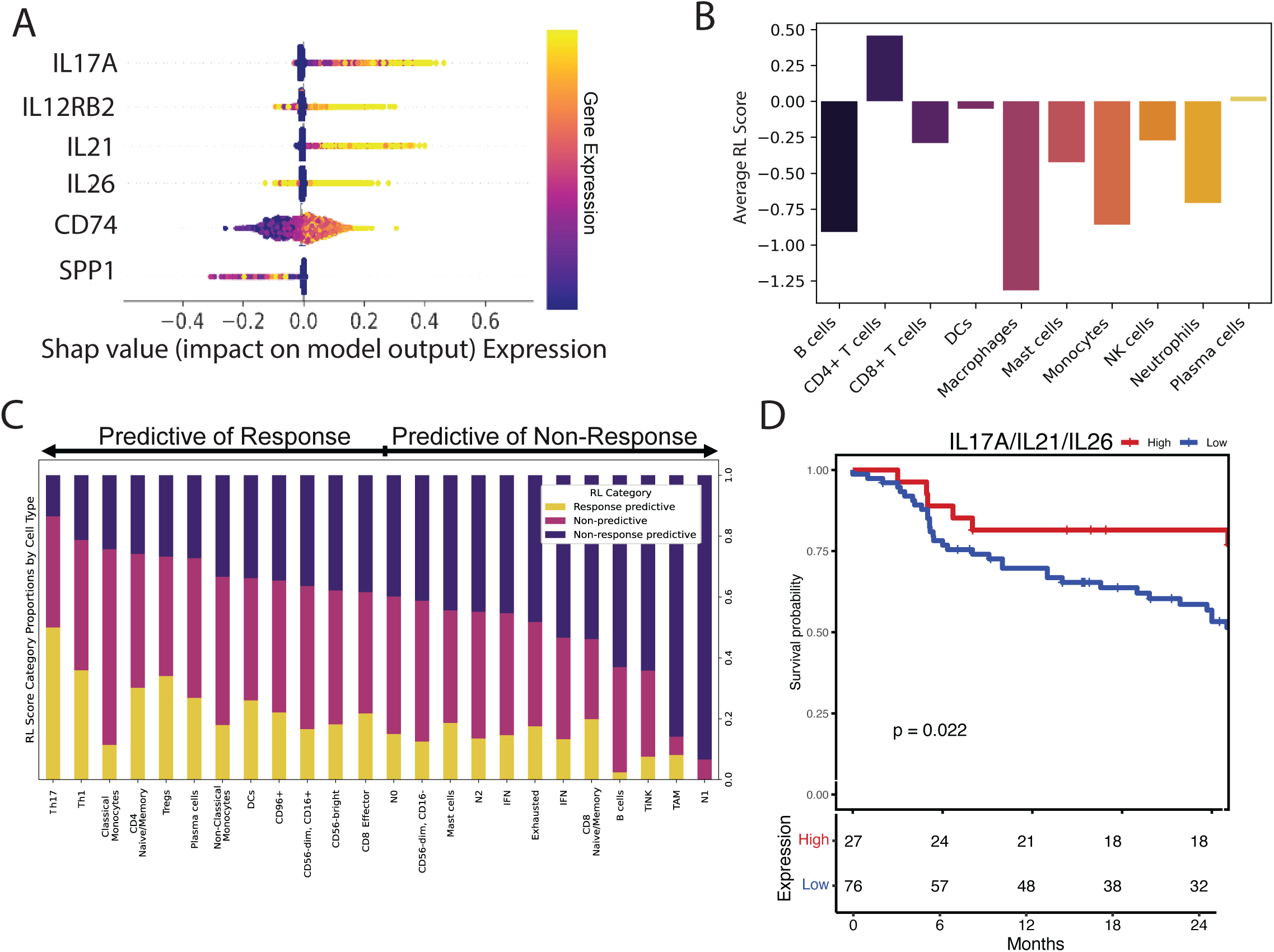
Machine learning model to infer important cellular and gene characteristics of BCG responders. **a.** Representative Boruta selected genes. Positive SHAP values indicate that gene is predicative of BCG responders. **b.** Reinforcement learning (RL) score averaged for each major cell type. Positive RL scores indicate the cell type is predictive of BCG responders. **c.** Proportion of a cell subtype’s reinforcement learning score (RL) categorized by response predictive (>0.5), non-response predictive (< 0.5), or non-predictive (−0.5, 0.5). **d.** Kaplan-Meier curve of recurrence-free survival for patients expressing high and low expression of IL21, IL17A, and IL26 defined with a log-rank p-value.

To determine which immune cell types are most critical for predicting therapeutic outcomes, we applied Precise’s reinforcement learning framework using the Boruta-selected gene list. In this approach, initial cell labels were assigned as +1 for cells from BCG responders and −1 for those from non-responders. These labels were then iteratively updated based on the model’s accuracy in classifying each cell, allowing us to quantify the predictive contribution of each cell type. Remarkably, CD4+ T cells emerged as the only population with a net positive predictive value for BCG response, reinforcing their unique role in driving effective anti-tumor immunity (Figure 5B). In contrast, while multiple cell types contributed to predicting non-response, macrophages stood out as the strongest indicators of treatment failure (Figure 5B). This dichotomy suggests that an effective BCG response hinges on the presence of immunostimulatory CD4+ T cell subsets, whereas an immunosuppressive myeloid environment dominated by macrophages may underlie therapeutic resistance.

To better understand which immune populations are associated with BCG treatment outcomes, we evaluated each cell subtype based on categorizations of the reinforcement learning scores. We assigned a classification to each cell individually with a cell scoring less than –0.5 as “predictive of non-response” and a score greater than +0.5 as “predictive of response.” Cells with scores near zero indicating no predictive power and termed “non-predictive.” Ranking cell types by these categorizations revealed patterns consistent with known biology (Figure 5C). For instance, exhausted CD8+ T cells were more strongly linked to BCG non-response, whereas effector CD8+ T cells were associated with response (Figure 5C). Similarly, pro-inflammatory classical monocytes and cytotoxic CD16+ NK cells had higher predictive scores than their immunosuppressive counterparts, such as tumor-infiltrating NK cells (TiNKs) and TAMs (Figure 5C). Notably, Th17-like Th1 cells emerged as the most predictive subset for BCG responsiveness (Figure 5C). Together, these results demonstrate the capacity to Together, these results underscore how the overall immune cell composition shapes the trajectory of BCG treatment response.

Given that Th17-like Th1 CD4+ T cells were the most enriched population in BCG responders, and that IL-17A, IL-21, and IL-26 emerged as top predictive features in our machine learning analysis, we sought to validate this observation in an independent dataset. Using our previously published cohort of 103 BCG-treated patients, we stratified individuals based on their cytokine expression. Strikingly, patients with high expression demonstrated significantly improved recurrence-free survival over 24 months, with 18 out of 27 (66%) remaining recurrence-free, compared to only 32 out of 76 (42%) in the low expression group (Figure 5D). These results suggest that a robust Th17-like Th1 response may play a pivotal and previously underappreciated role in orchestrating effective anti-tumor immunity in the context of BCG therapy.

## Discussion

In this study, we define the immune landscape of the bladder during intravesical BCG therapy using single-cell transcriptomic profiling of bladder-infiltrating and circulating immune cells from patients with high-risk NMIBC. By integrating cell state analysis, cell–cell communication, and machine learning, we identify distinct immune programs that segregate BCG-responsive from non-responsive patients. Our data highlight a Th17-like Th1 CD4+ T cell subset, robust antigen presentation, and reduced immunosuppressive signaling as hallmarks of BCG responsiveness. In contrast, BCG unresponsiveness is characterized by regulatory T cell (Treg) dominance, exhausted CD8+ T cells, and macrophage-driven immunosuppression.

While the exact immune-driven mechanism of BCG in bladder cancer is not fully understood, it likely follows a similar path to the conventional mechanism underlying control of mycobacterial infections such as tuberculosis. Here, it is understood that mycobacteria are taken up by macrophages, where antigen presentation through MHC molecules and secretion of IL-12 promote differentiation of Th1 CD4+ T cells. These cells produce IFN-γ and TNF, which promote the formation of granulomas, organized structures of chronically activated macrophages encircled by lymphocytes that continuously preserve granuloma integrity. In *Mtb*, such immune complexes serve to contain mycobacteria in a latent state, and in bladder cancer, they likely function as a localized hub of sustained immune activation. Supporting this, histological studies of BCG-treated tissue have found that granuloma formation correlates with improved recurrence free survival^39,40^.

BCG treatment is known to increase circulating monocytes and drive macrophage infiltration into the bladder wall and are elevated in the urine^41^. Clinically, a higher density of CD68+ macrophages in tumors following intravesical BCG immunotherapy correlates with improved recurrence-free survival in NMIBC patients^41^.

Paradoxically, however, elevated numbers of TAMs prior to treatment are linked to poor outcomes, suggesting that macrophage origin and polarization state critically shape therapeutic efficacy^41^. As evidenced by *Mtb* infections, granuloma formation is primarily sustained by M1-polarized macrophages in latent disease, whereas M2 polarization correlates with an uncontrolled active disease state^42^. By analogy, resident TAMs in the bladder tumor microenvironment are thought to be primarily in an M2 immunosuppressive state, whereas newly recruited monocytes differentiate into more activated M1-like macrophages that promote tumor control^31,32^. Our data support this model, as we observed that BCG responders not only had more activated gene signatures in TAMs, but also had significantly higher frequencies of classical monocytes in their blood.

These classical monocytes are known to differentiate into M1-polarized macrophages, whereas non-classical monocytes give rise to M2-like macrophages^31,32^. This increase in classical monocytes, together with the reduction in macrophage immunosuppression signature in BCG responders, suggests that recruitment and differentiation of fresh monocytes into inflammatory macrophages is critical for anti-tumor activity. Consistent with this notion, studies in B16 melanoma have shown that intravenous BCG induces monocyte populations that differentiate into inflammatory macrophages essential for tumor control, and transfer of BCG-conditioned bone marrow is sufficient to protect against tumor growth^43^. Together, these findings highlight monocyte-derived macrophages as key mediators of in determining effective BCG immunotherapy outcomes.

Macrophages then orchestrate the adaptive immune responses by presenting antigen and shaping the differentiation of CD4+ and CD8+ T cells through cell-cell interactions. We found that BCG non-responding TAMs had increased immune checkpoint inhibitor interactions with T cells through PDL1, PDL2, TGFb, CD48, and TRAIL. Since such markers have been found to influence the differentiation of T cells, these interactions suggest that macrophages contribute to the exhausted lymphocyte profile of BCG-unresponsive patients, which is characterized by a higher proportion of exhausted CD8+ T cells and Tregs. Rather than expression by the bladder epithelium, this suggests that bladder resident macrophages could be a source of exhaustion after BCG.

By comparison, the macrophages in BCG responders had a less suppressive phenotype, with markedly higher expression of MHC II genes. Since *Mtb* and BCG have a primarily phagosomal lifecycle, adaptive immune identification is chiefly mediated through presentation of peptide:MHCII^44^. This emphasizes the likely importance of the CD4+ T cell compartment in BCG response, and why HIV infection is such a strong risk factor for tuberculosis^45^. A key observation we found was the emergence of Th17-like Th1 CD4+ T cells in BCG-responsive patients that underwent potent expansion. These cells expressed both canonical Th1 molecules (e.g., IFNG, STAT4, TBX21) and Th17-associated cytokines (IL17A, IL17F, IL26, IL21). Due to the low number of macrophages, we could not comprehensively examine lowly expressed cytokines that likely play a role in this differentiation process, such as IL-12 or IL-6. While we could not identify a definitive interaction with macrophages that was implicated in this differentiation, prior studies in other Mycobacterial models have found that M1-like macrophages induce Th1 and Th17 cell responses^46^. Together, this suggests pro-inflammatory macrophages in BCG responders could be playing a key role in this differentiation.

In addition, we found that these Th17-like Th1 cells were not only more abundant in responders, but also were the gene signature enriched in our machine learning model, suggesting they represent a qualitatively superior population in mounting local anti-tumor immunity. In particular, the cytokines IL-21, IL-26, and IL-17A were identified as the most predictive for BCG responsiveness, which was subsequently validated in a separate cohort. These genes have several complex and conflicting roles; for example, IL17A has been correlated with higher-grade tumors and is known to promote angiogenesis^47^. On the other hand, early expression of IL17A is critical for recruitment of neutrophils^48^. Alternatively, Th17 cells are known to compete with Tregs for TGF-β, so the generation of Th17 cells could simply be a way to prevent the development of immunosuppressive Tregs^22^. While this Th17-like Th1 cell’s role in BCG response is not completely understood, studies in *Mycobacterium leprae* have found that Th17 gene signatures are correlated with reduced *M. leprae* burden^49^. Finally, we found that these Th17-like Th1 cells produced higher levels of IL-21 and CCL5 (also known as RANTES). These cytokines exemplify classic CD4+ T cell “help,” whereby CD4+ T cells provide supportive signals, through cytokines and co-stimulatory interactions, that enhance CD8+ T cell activation, proliferation, and effector differentiation^24,37^. Our cell–cell interaction analysis, together with prior literature, suggests that IL-21 and RANTES from Th17-like Th1 cells contribute to the priming and functional maturation of CD8+ T cells into cytotoxic effector cells.

Our work identifies the heterogeneity and immune landscape associated with responses to BCG in non-muscle invasive bladder cancer. We identify critical changes in T cell and myeloid cell populations are associated with clinical resistance to BCG and identify critical population changes and cellular interactions of interest for future investigation. Our results identify that there are parallel signaling pathways between persistent *Mtb* infection of the lung and resistance to BCG therapy. Future studies that leverage our identified population changes and cellular interactions may be applied to overcome BCG resistance and treatment response by risk stratification and combinatorial therapies.

Finally, we acknowledge limitations to these findings. While this investigation is one of the largest single-cell analysis of BCG treated bladders, an increased number of patients would potentially refine subset differences in cell types that had a limited number of cells and remove any batch effects identified in our machine learning model. Moreover, our cell-cell interactions were inferenced based on predicted interactions, future investigations of multi-dimensional omics could validate the three-dimensional nature of immune cell interactions.

## Supplementary Figure Legends

**Supplementary Figure 1. Immune cell gene expression and distributions validation**

**a.** Feature plot of representative genes for each major cluster.

**b.** UMAP plot of cells in both the blood and tissue.

**c.** Stacked bar plots summarizing cell subset proportions after down sampling to 200 cells per sample in the blood tissue.

**Supplementary Figure 2. Distribution of immune cell types across blood and tissue**

**a.** Stacked bar plot summarizing cell count (top) and cell proportions (bottom) for each patient’s bladder.

**b.** Stacked bar plot summarizing cell count (top) and cell proportions (bottom) for each patient’s blood.

**Supplementary Figure 3. Cytotoxic and effector NK cell signatures are enriched in BCG responders**

**a.** UMAP of subset NK cells from the blood and bladder tissue.

**b.** Dot plot of representative genes expressed in each major cluster. Dot size represents percent of cells expressing the gene; color represents scaled expression of the gene.

**c.** Slingshot trajectory of NK T cells.

**d.** Correlation between a patient’s proportion of CD56-dim, CD16+ cells and TrNK cells.

**e.** Proportion of NK cell subsets in blood and bladder. Box plots showing the distribution of immune cell type proportions for each patient.

**f.** Proportion NK cell subsets in BCG naïve, responsive, and unresponsive groups in the bladder tissue. Stacked bar plots summarizing NK cell subset proportions after down sampling to 50 cells per sample.

**g.** Proportion of NK cell subsets in BCG naïve, responsive and unresponsive groups in the bladder tissue.

**h.** Proportion of NK cell subsets in BCG naïve, responsive and unresponsive groups in the blood.

**Supplementary Figure 4. CD4+ T cell composition in blood and bladder**

**a.** Cosine similarity scores between subsets of CD4+ T cells.

**b.** UMAP of blood and bladder tissue cells.

**c.** Proportion of CD4+ cell subsets in BCG naïve, responsive and unresponsive groups in the blood.

**Supplementary Figure 5. CD8+ T cell composition in blood and bladder**

**a.** Feature plot of representative genes for each major cluster.

**b.** UMAP plot of cells in both the blood and tissue.

**c.** Stacked bar plots summarizing cell subset proportions after down sampling to 50 cells per sample in the blood tissue.

**Supplementary Figure 6. BD-Rhapsody identifies cellular diversity of neutrophils responding to bladder cancer.**

**a.** UMAP of neutrophils from the blood

**b.** UMAP of neutrophils from the bladder tissue

**c.** Dot plot of representative genes expressed in each major cluster. Dot size represents percent of cells expressing the gene; color represents scaled expression of the gene.

**d.** Proportion of neutrophils cell subsets in blood and bladder. Box plots showing the distribution of immune cell type proportions for each patient.

**e.** Proportion of neutrophils cell subsets in BCG naïve, responsive and unresponsive groups in the blood.

**f.** Proportion of neutrophils cell subsets in BCG naïve, responsive and unresponsive groups in the bladder tissue.

**g.** Proportion of neutrophils cell subsets in BCG naïve, responsive and unresponsive groups in the bladder tissue when including all other cell types.

**h.** Proportion neutrophils cell subsets in BCG naïve, responsive, and unresponsive groups in the bladder tissue. Stacked bar plots summarizing neutrophils cell subset proportions after down sampling to 50 cells per sample.

**Supplementary Figure 7. Monocytes and macrophage composition in blood and bladder**

**a.** UMAP of monocyte and macrophage subsets between the blood and bladder tissue.

**b.** Proportion of monocyte and macrophage cell subsets in BCG naïve, responsive and unresponsive groups in the blood.

**c.** Proportion of monocyte and macrophage cell subsets in BCG naïve, responsive and unresponsive groups in the bladder tissue.

**Supplementary Figure 8. CD4+ T cell interactions define responders**

**a.** Dot Plot of MHCII cellular interactions between antigen presenting cells and CD4+ T cells.

**b.** Survival curve of CD74 and HLA-DOA over the course of two years.

**c.** Stacked bar plot of signaling found between CD4+ T cells and CD8+ T cells in BCG responders and non-responders (left). Expression of CCL5 in CD4+ T cells (middle) and CCR5 in CD8+ T cells (right).

**Supplementary Figure 9. Machine learning identifies responder and non-responder gene sets.**

**a.** Dot Plot of Boruta selected genes grouped by cell type expression

**b.** SHAP value scores for Boruta selected genes.

## Methods

### Sex as a biological variable

Male and female participants were included in the study; however, patients were included based on availability and not did not end up evenly across conditions. Women were only in the BCG naïve and unresponsive groups; thus it is unknown if the BCG responsive findings are relevant for women.

### Sample Processing

Tissue was transported in DMEM with 10% FBS to preserve stability at room temperature to preserve temperature sensitive granulocytes. Tissue was minced and digested in two rounds (20 minutes and 15 minutes) in DMEM without FBS using Liberase TM (Sigma) in a 37 °C water bath with gentle manual inversion. The digested tissue was resuspended in FACs buffer passed through a 70 uM strainer (Corning), centrifuged and subjected to CD45 enrichment using the Miltenyi biotech CD45 microbeads as per the manufacturer’s instructions.

In parallel, 1 ml of blood from blood isolated in EDTA coated tubes was also processed to isolate leukocytes using whole blood lysis protocol recommended by 10X Genomics (CG000392) using eBioscience 1X RBC Lysis Buffer (Thermo fisher) with PBS washes and low speed centrifugation.

### BD Rhapsody single-cell RNA sequencing

To combine the matched tissue and blood samples to load on a single BD Rhapsody cartridge, the processed matched tissue and blood samples were labeled with sample tags using the BD Single-cell Multiplexing kit in accordance with the manufacturer’s protocol. The individual samples were resuspended in BD Rhapsody sample buffer (BD Biosciences) and stained with DRAQ7 and Calcein AM florescent dyes to enable visualization for counting and calculation of quality control metrics in the BD Rhapsody single cell system. The cells were counted, and blood-tissue samples mixed, passed through a 70 micron strainer and up to 10,000 cells loaded in the BD Rhapsody microwell cartridge system. Magnetic capture beads were then loaded close to saturation so that each microwell contains one magnetic bead. Each of the cells were isolated in the microwells, lysed and the polyadenylated mRNA molecules from each cell captured by individual magnetic capture beads containing unique molecular identifiers (UMIs) and cell specific barcodes. The capture beads were retrieved from the BD Rhapsody single cell system and cDNA reverse transcription carried out with cDNA with UMI and cell specific barcode information added to the cDNA molecules. 3 ’end whole transcriptome amplification (WTA), sample tag libraries were prepared according to the BD Rhapsody WTA and sample tag preparation protocol. For quantification and quality control purposes the libraries were subjected to Qubit and Tape station analyses. For calculation of total paired end reads, sequencing depth of 60,000 reads per cell were used for the WTA library analyses, for the sample tag libraries sequencing depth 1200 reads per cell. The WTA and sample tag libraries were mixed according to custom BD Rhapsody calculations. The libraries were subjected to sequencing using 150 bp paired end sequencing using the Novaseq X Plus by Admera Health (New Jersey).

### Data Processing

The BD Rhapsody fastq files were processed using the BD Rhapsody WTA pipeline, available on the Seven Bridges Genomics cloud server. Briefly the pipeline had steps which involved filtering of the reads, identifying the cell ID and the UMI, aligning the reads to the the hg38 reference genome or in the case of Abseq data, the Abseq reference file. The raw counts were error corrected, putative cells identified and the final processed file contained reads per cell for the samples generated along with the quality control metrics.

### Data filtering and quality control

The Seurat objects generated at the end of the BD Rhapsody WTA pipeline were downloaded and further processing was done using Seurat (v3) in R version 4.3.2/4.3.3^50–52^. Initially quality control filtering was carried out to remove low expressing cells or potential doublets as recommended by the Seurat pipeline removing cells with high percentage of mitochondrial reads and low expressing cells. Cells identified as having no sample tags (“Undetermined”) or identified as multiplets by the BD Rhapsody pipeline were also removed. To ensure that singlets were identified, DoubletFinder was run on each sample to remove doublets^53^.

### Data Analysis

All analyses were performed in R. Integration across all samples was performed with canonical correlation analysis. Analysis of cellular subsets were reintegrated using either SCTransform, CCA, or harmony. Determination of integration method was based on visualized batch effects on the umap plot. Data visualization was carried out using the ggpubr and ggplot2 packages. Box-and-whisker plots were generated with ggboxplot() specifying sample groups as factors, and statistical comparisons were overlaid using the stat_compare_means() function with appropriate tests (t test, Wilcoxon, or Kruskal–Wallis, depending on distribution). For box plots where percentages were calculated, a minimum cell threshold was set at 100 cells per patient to be included, and reduced by increments of 25 cells until each BCG condition or tissue type had at least 3 patients. Additional visualizations including stacked bar plots were generated with ggplot() using customized themes.

Trajectory inference was performed using the Slingshot package^54^. Input data consisted of dimensionally reduced coordinates from Seurat UMAP embeddings, along with cluster labels. Slingshot was run with default parameters, initializing trajectories from the earliest cell cluster, and pseudotime values were assigned to each cell based on principal curves fitted through the low-dimensional space.

For heatmap generation, we first identified differential expression between clusters through Seurat’s FindAllMarkers() function. Genes meeting significance thresholds (adjusted p < 0.05) were selected and visualized with the ComplexHeatmap package^55^. Heatmaps were constructed with Heatmap() using row scaling (scale = “row”) and Euclidean distance–based clustering. The number of cells per patient were down-sampled to remove differences in sequencing depth accross patients. Hierarchical clustering was initialized at the level of the number of predefined cell types, and the tree was iteratively cut into higher numbers of clusters until additional subdivisions resulted in redundant or overlapping gene modules.

Cell–cell communication was inferred with CellChat^56^. All cells from the bladder samples were included as input to createCellChat(). Normalization and preprocessing followed the CellChat workflow. Networks were inferred with computeCommunProb() using default parameters, and aggregated at the signaling pathway level with computeCommunProbPathway(). Communication probability matrices were then projected into signaling networks for visualization.

Machine learning analysis was conducted with the Precise package using the complete bladder dataset as input. Gene selection was first performed using the Boruta algorithm. The resulting gene set was used to train an XGBoost model. The final model was integrated with Precise’s reinforcement learning framework, which iteratively updates predictive weights based on misclassified cells. Each cell was assigned a score corresponding to its predicted class probability, which was used in downstream analyses.

### Statistics

Statistical comparisons for sequencing data were performed in R. Comparisons of between blood and tissue compartments was performed with the stat_compare_means function using the Wilcox test, while comparisons between BCG naïve, responsive, and unresponsive groups was performed with the Kruskal Wallis test.

Comparisons of cellular distributions in stacked bar plots was compared through a chi-square test. Linear regressions were performed using the lm function.

### Study Approval

Patients were prospectively under IRB approved clinical research STU00219216. Peripheral blood and tissue were collected from the patients at the time of surgery and immediately for standard of care bladder cancer resection and processed for single cell RNA sequencing. Sample size was not determined prior to the prospective collection of samples and response to BCG was determined by the pathology report post-surgery. Informed consent was received prior to participation.

## Data Availability

Once published, all code will be made available raw sequencing will be made available on EGA. The original R and Python code used to generate most Figure panels will be publicly available as of the publication date. Any additional code may be requested from the corresponding author.

## Author contributions

Conceptualization, MK, JJM, Methodology: RJB, MK, JP, WC, JQ, YY, SK, LL, CC CMW, BC, JJM Software: RJB, MK, Validation, RJB, MK, WC, BC, JP Formal Analysis, JJM, RJB, HN, MK, JP, BC. Investigation, All authors Resources, CMW, SK, LL, CC. Data Curation, All authors Writing: Original Draft, RJB, MK, JJM, JP. Writing: Review & Editing: All authors, Visualization: RJB, MK, JJM, JP. Supervision, JM, JP, BC, WC Project Administration, YY, SK, LL, CC. Funding Acquisition: JJM

## Acknowledgments

We sincerely appreciate the enrollment of clinical coordinators Lydia Landrum, Sophia Kallas, Claire Carter. The authors would like to thank Jun Qian and Yanni Yu for help with blood processing. The authors would like to thank Khyati Meghani for useful discussions during the study.

“The project described was supported by the Robert H. Lurie Comprehensive Cancer Center

## H Foundation Core Facility Pilot Project Award to MK

The authors would like to thank Nicholas DePatie, Chloe Charendoff, Ashley Walton and Matthew Chang from BD BioSciences for technical support related to BD Rhapsody experimental running and bioinformatic analyses. JJM is funded by VHA BX005599 and BX003692 awards from the VHA.

