## Supplementary figures and images for "Divergent Macrophage-Regulated T cell States Determine Response to Bacillus Calmette-Guérin (BCG) in High-Risk Bladder Cancer"

### Supplemental Figures

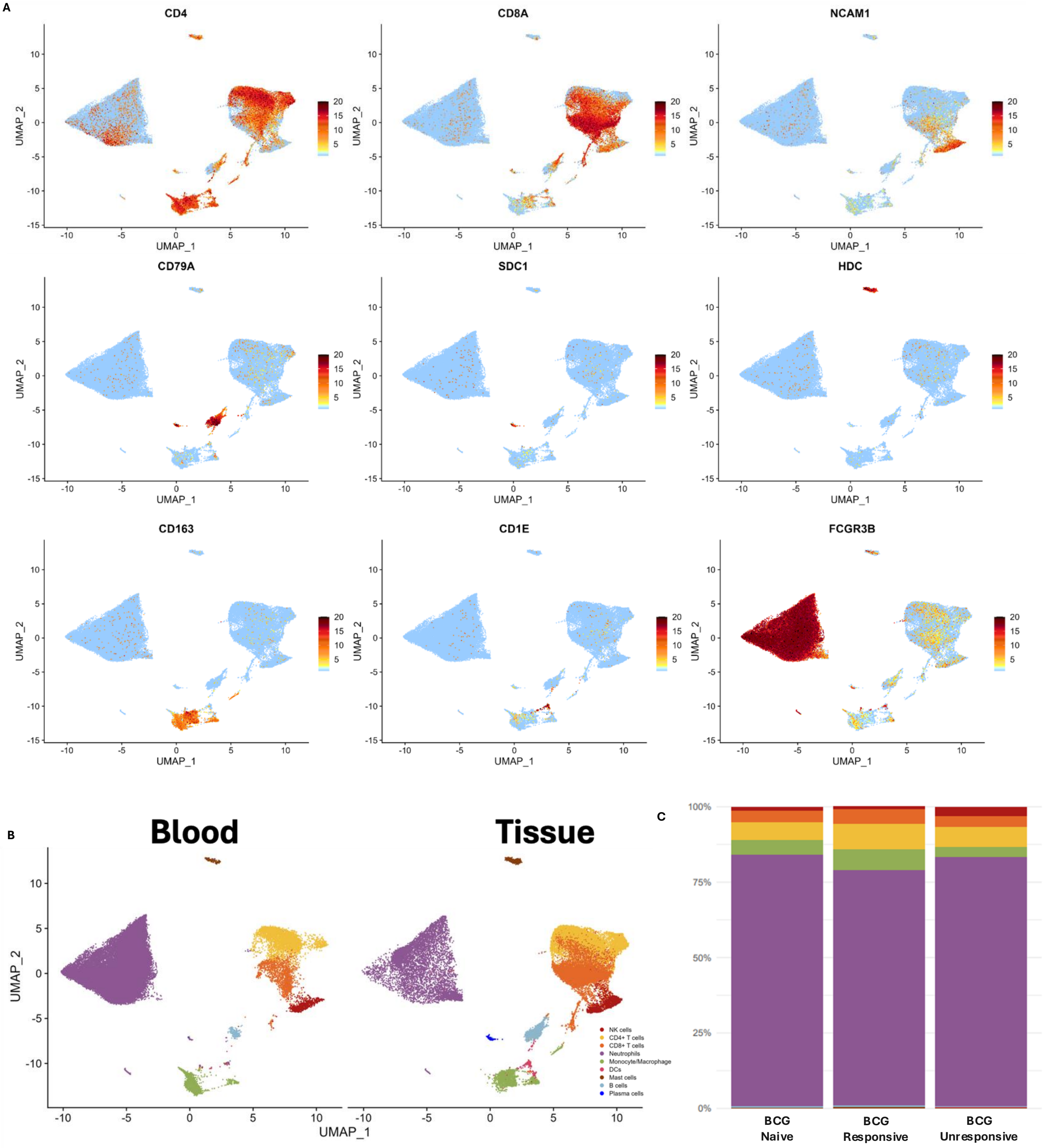

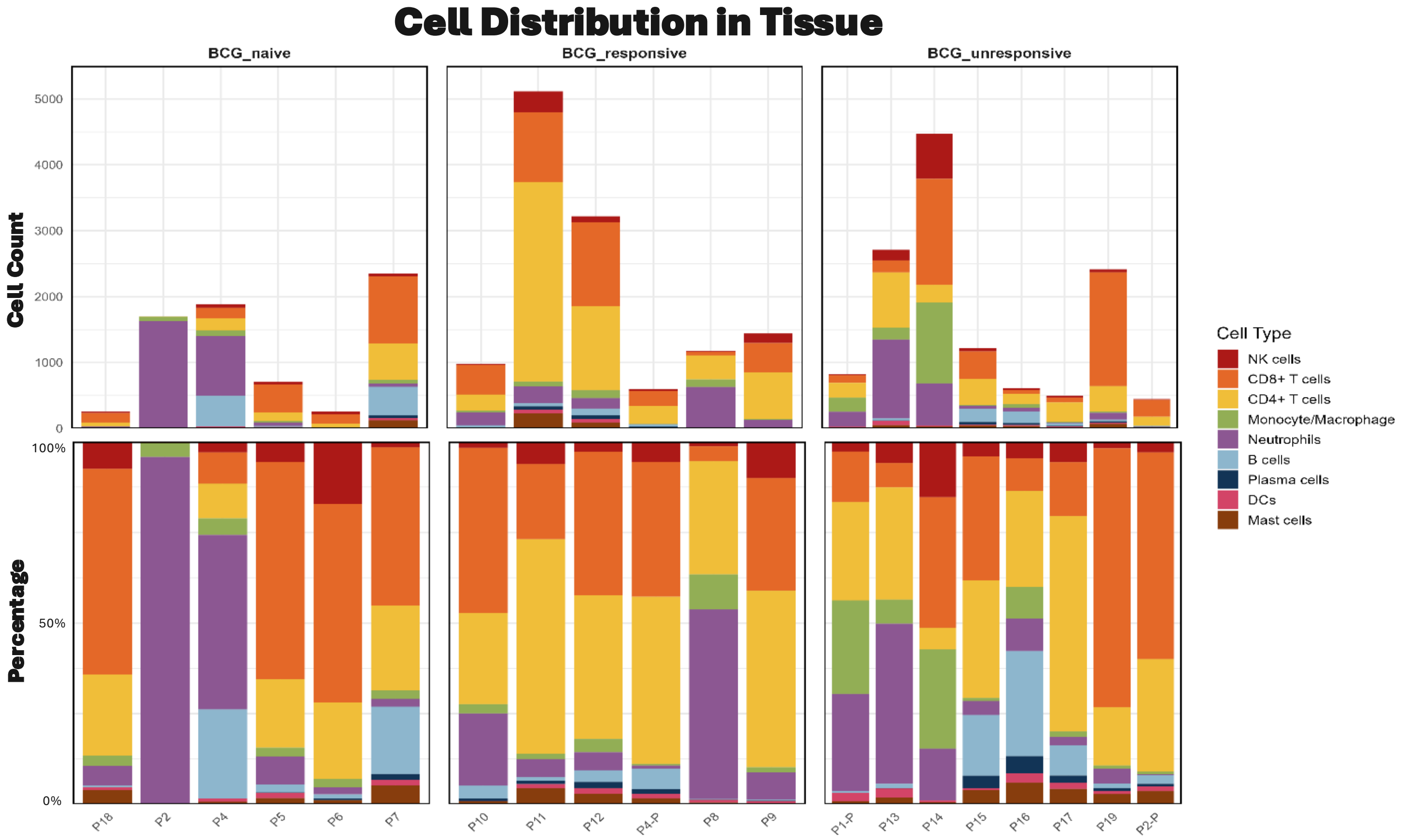

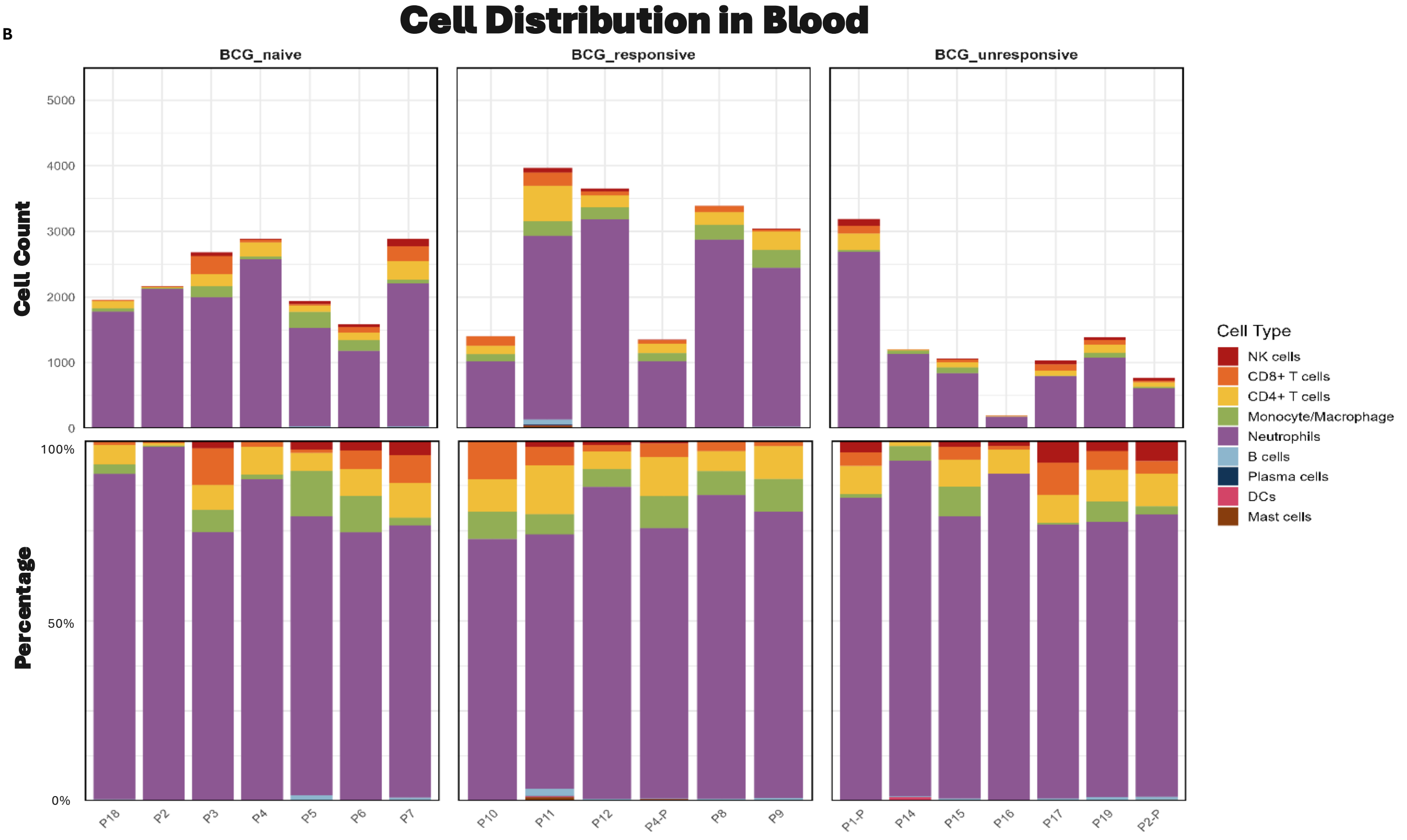

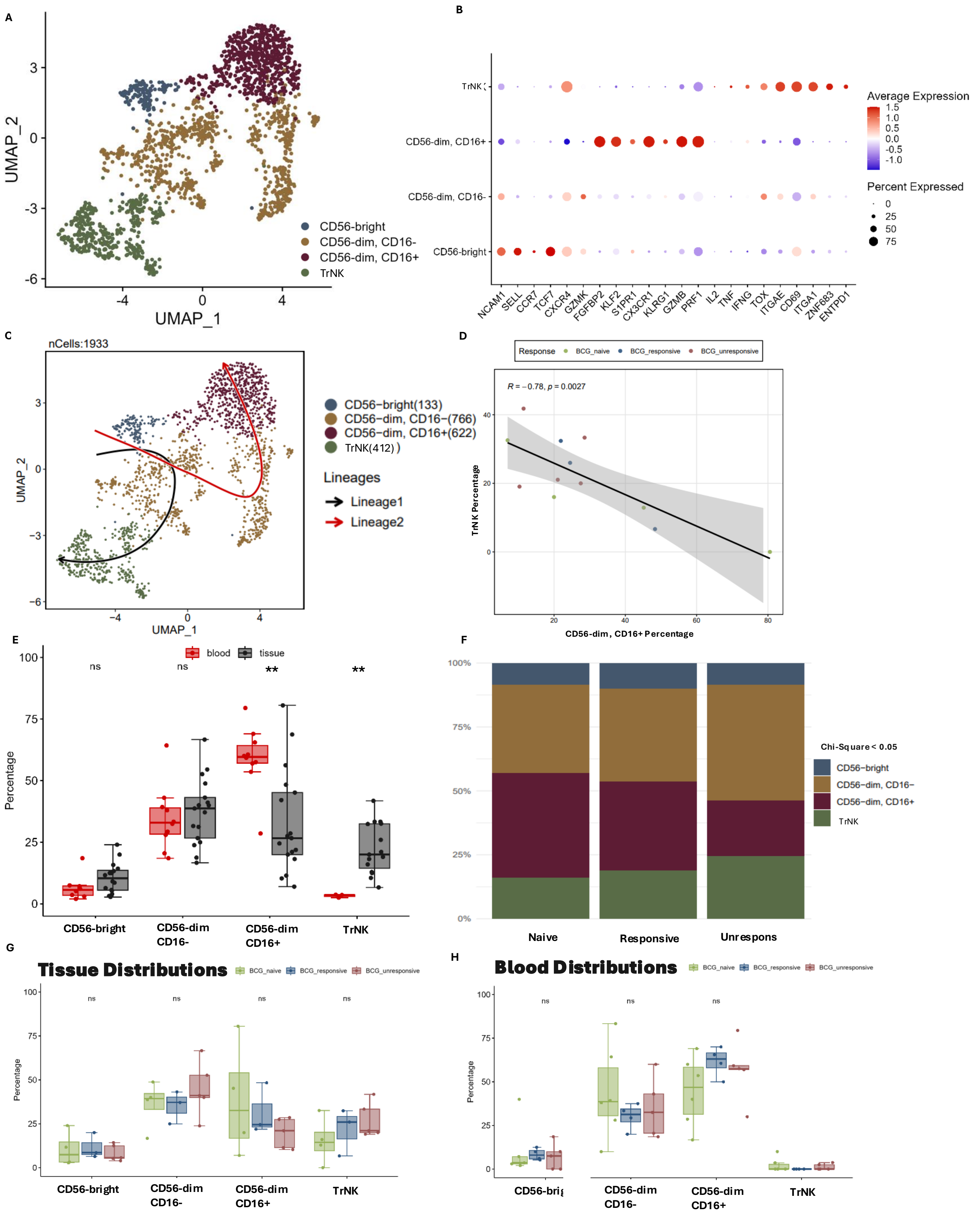

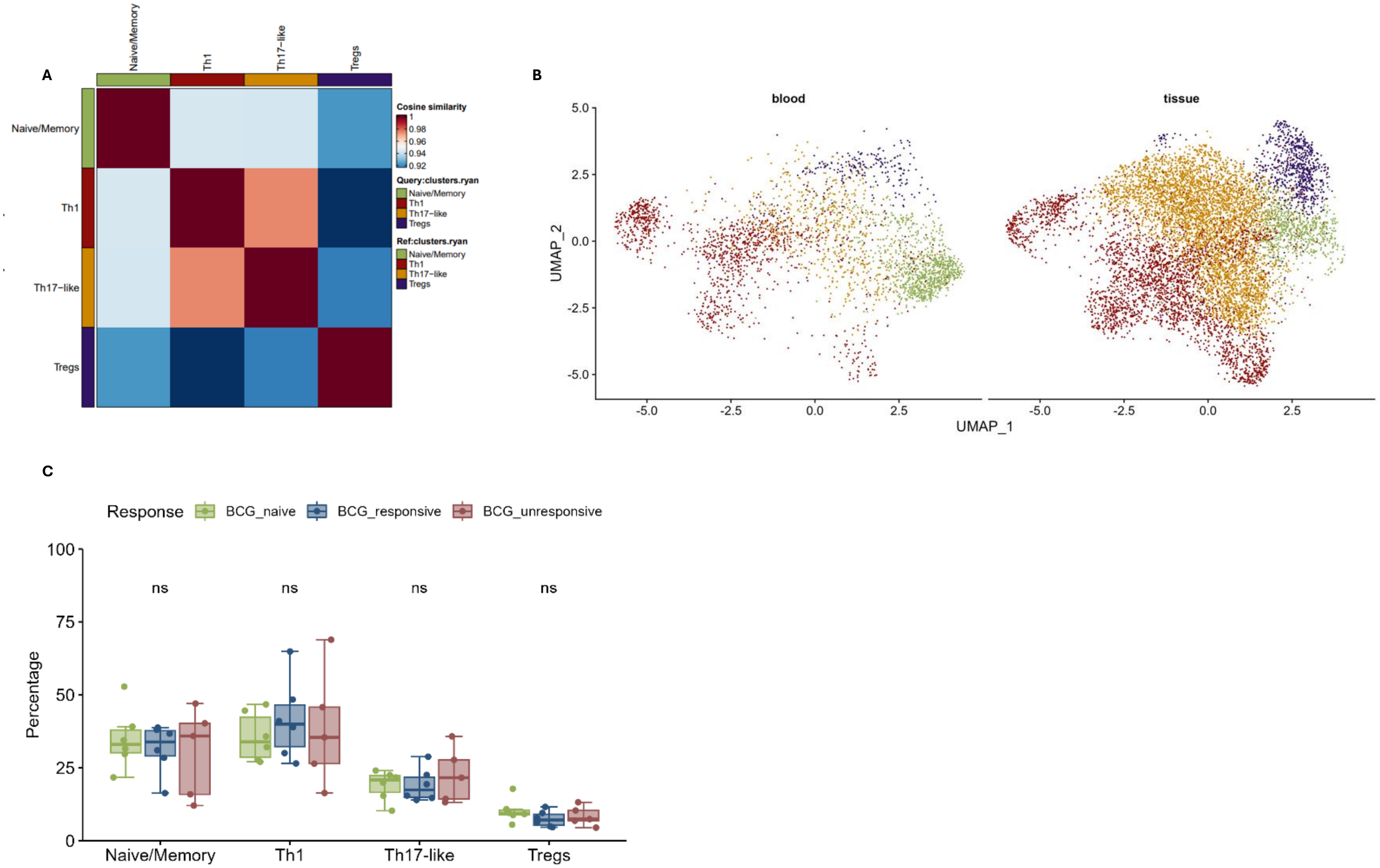

Figure S5

A

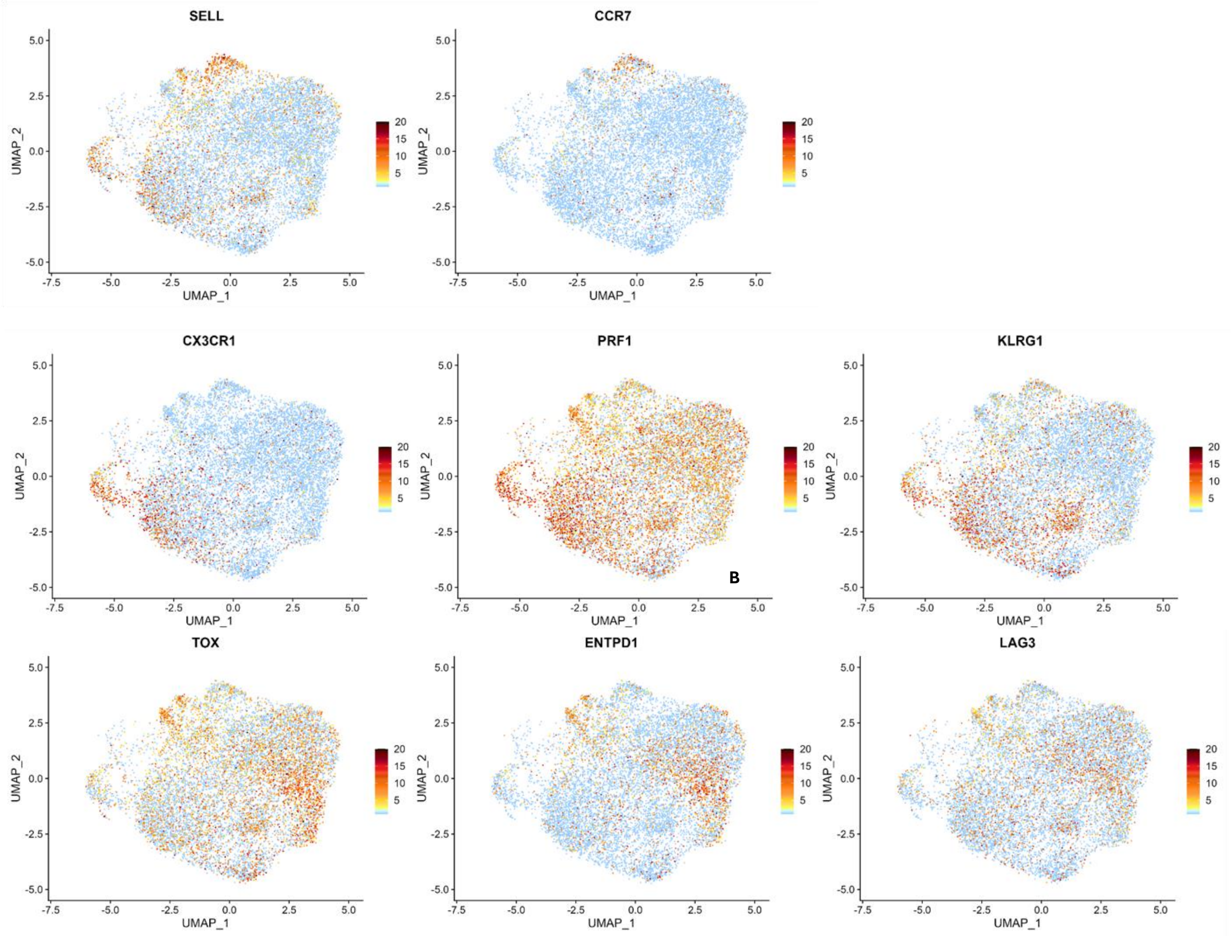


B

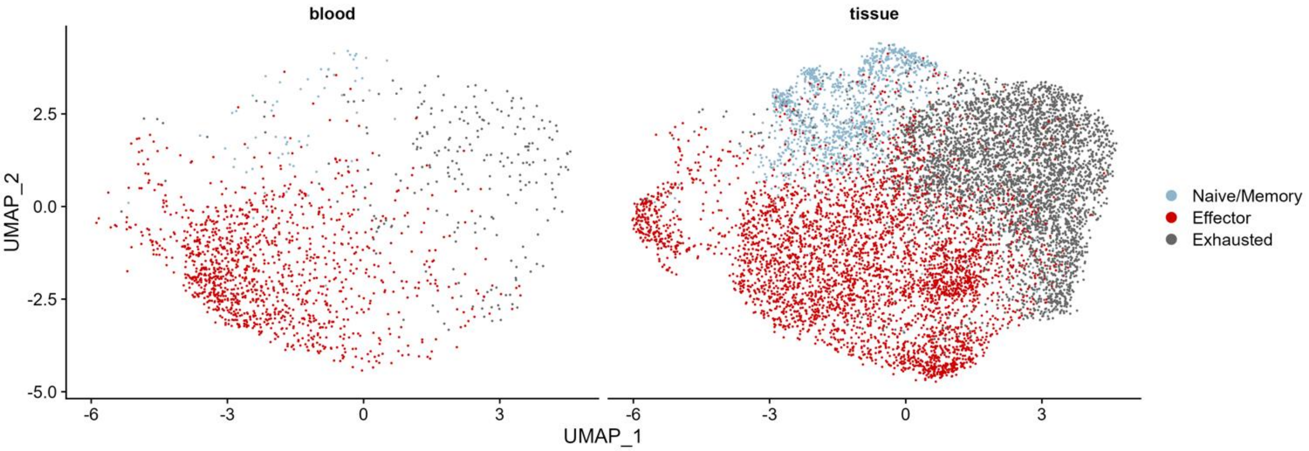


C

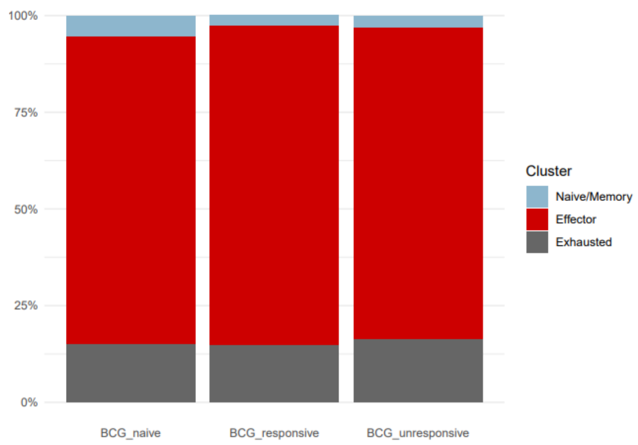

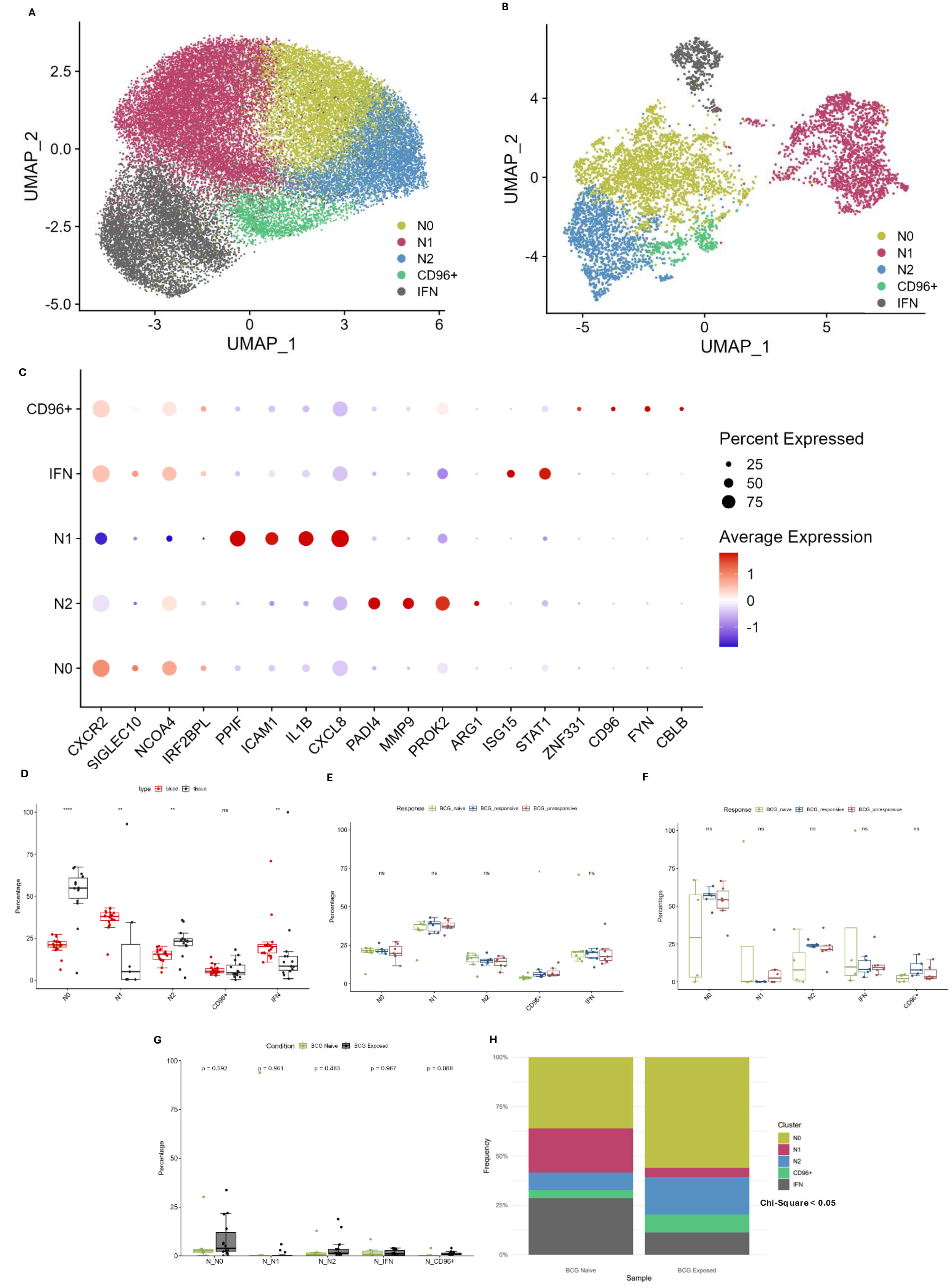

Figure S7

A

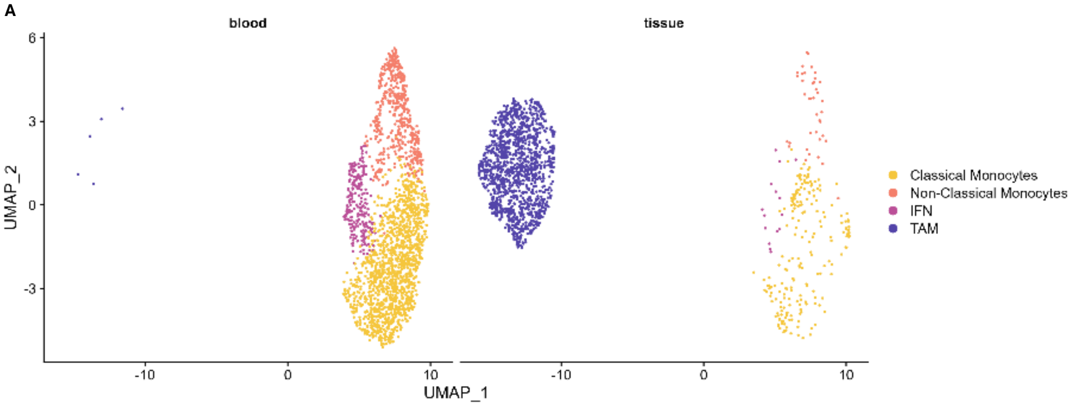


B

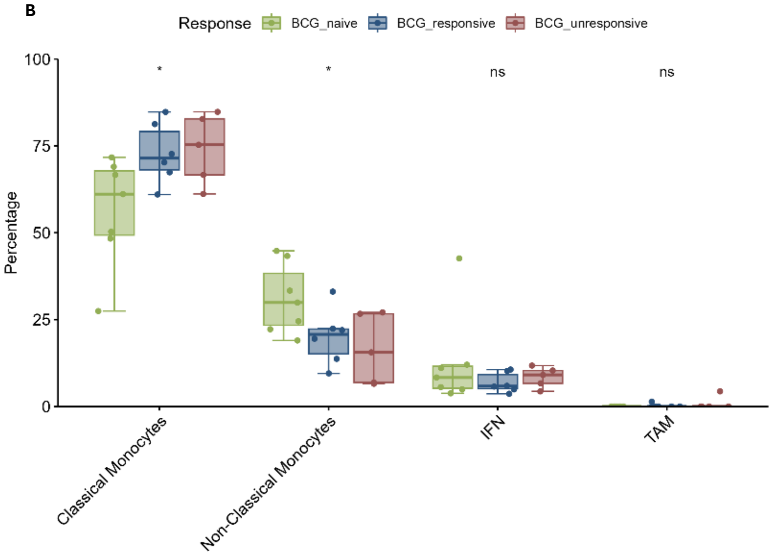


C

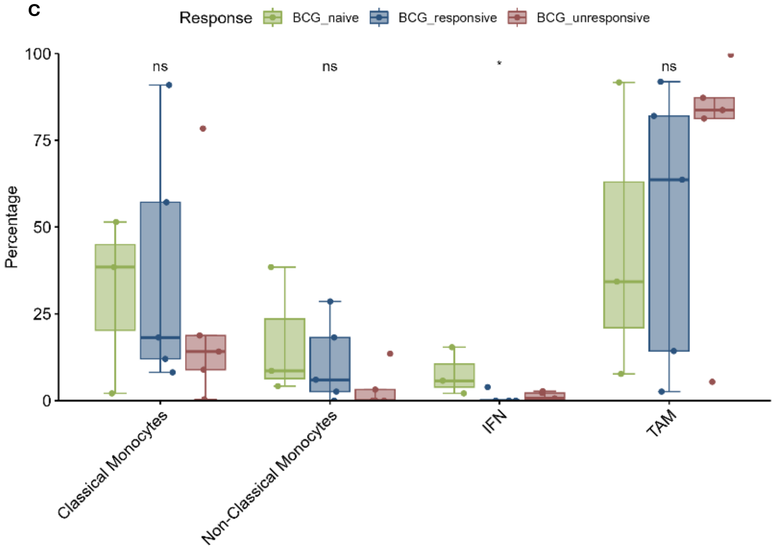


D

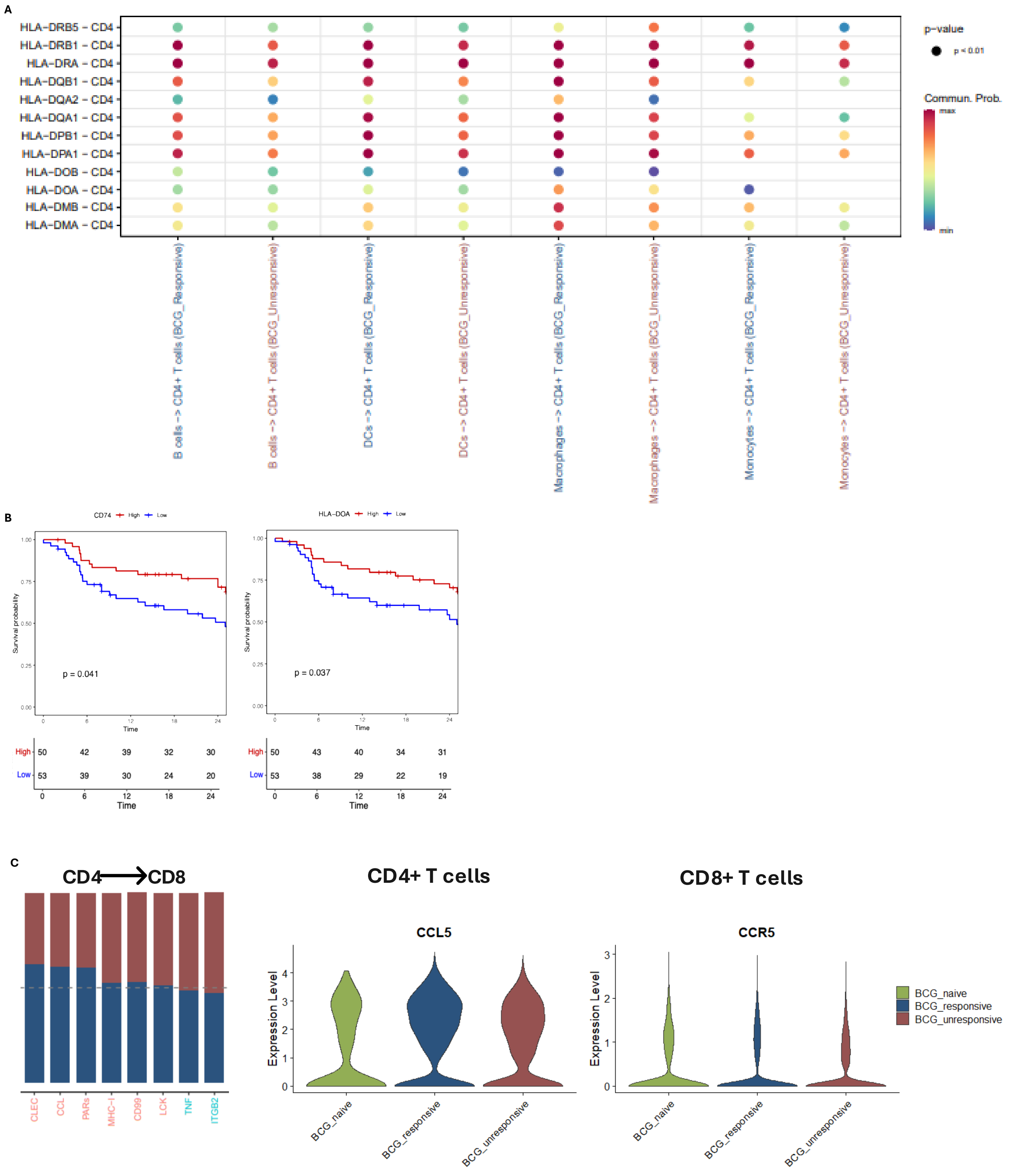

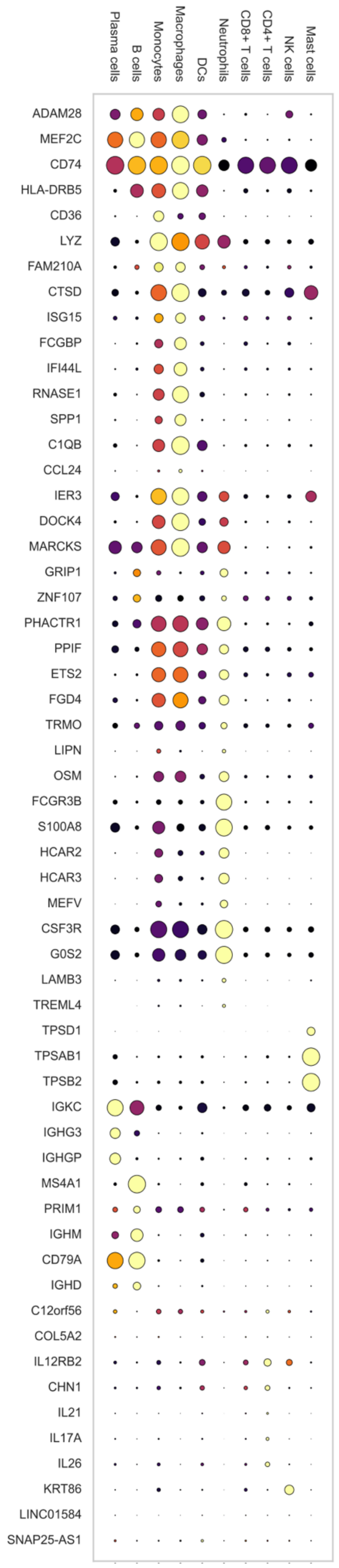

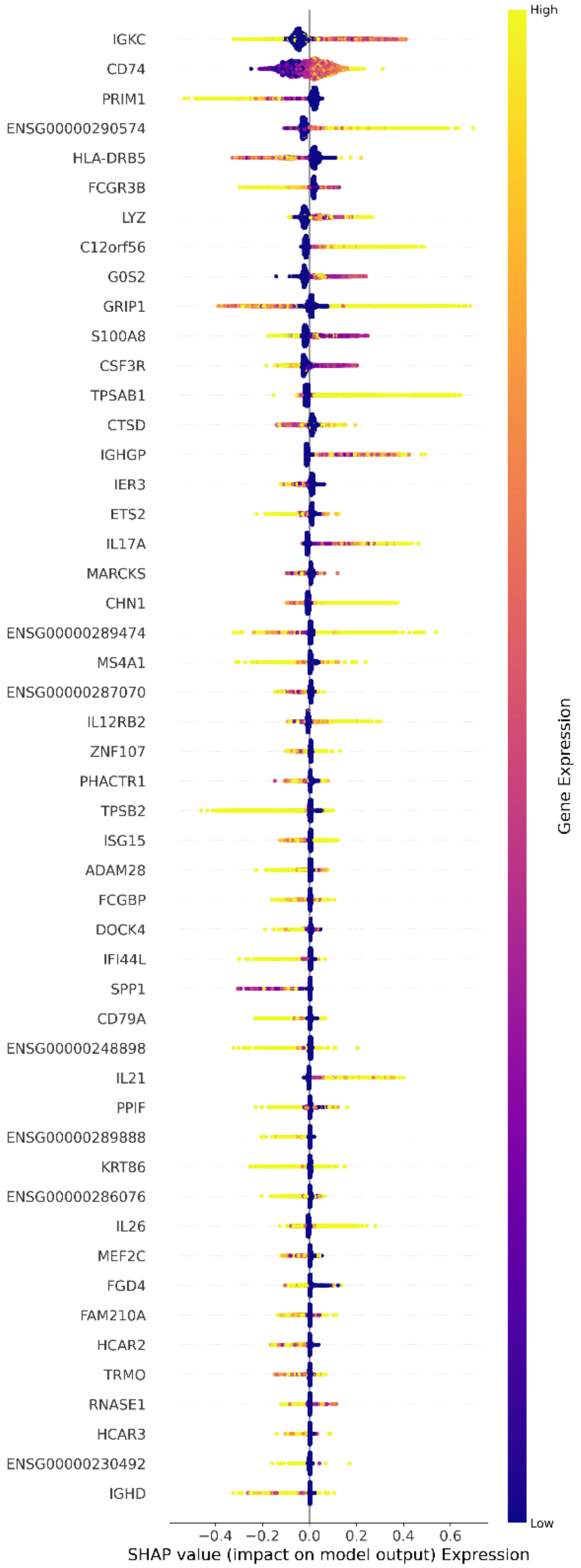
